# Focused ultrasound alters neural time irreversibility independently of response amplitude

**DOI:** 10.64898/2026.04.26.720929

**Authors:** Jacek P. Dmochowski, David H. Wolpert

## Abstract

Transcranial focused ultrasound (tFUS) can modulate activity in deep brain regions non-invasively, yet its effects remain variable and incompletely understood. Because sonication deposits mechanical energy into tissue, we asked whether responses could be clarified through the lens of stochastic thermodynamics, which links nonequilibrium dynamics to dissipation and time-irreversibility. Analyzing previously published fiber-photometry recordings from freely moving mice, we estimated the entropy production rate (EPR) before, during, and after thalamic sonication. GCaMP fluorescence rose monotonically with acoustic dose, but EPR followed a non-monotonic profile peaking at moderate intensity. Immediately after sonication, EPR was elevated yet uncorrelated with GCaMP, and retained a significant dose relationship even after accounting for response amplitude. Finally, trials with lower baseline EPR showed larger subsequent changes in both EPR and GCaMP during sonication. These findings indicate that tFUS reshapes neural dynamics beyond what signal amplitude captures, while highlighting the insights afforded by adopting stochastic thermodynamics in neuromodulation research.

## 1 Introduction

Transcranial focused ultrasound (tFUS) has emerged as a promising non-invasive approach for modulating deep brain activity with millimetre-scale spatial precision and access to targets that are difficult to reach with electromagnetic stimulation [5, 6, 17, 27, 38, 49, 50]. Realization of this promise will require addressing the widely observed variability in the direction and magnitude of the brain’s response to stimulation across subjects, sessions, and trials [7, 14, 36, 37, 39, 51, 56]. Potential sources of this variability include biophysical differences between subjects [14], differences in experimental design across studies [37, 56], and the inherently non-stationary nature of neural activity – stimulation interacts with ongoing dynamics, such that the response is itself modulated by the baseline state [39]. In the transcranial magnetic stimulation (TMS) literature, state-dependent effects are well documented [4, 48]. Such findings suggest that explaining response variability will require modeling tFUS as a perturbation of a non-stationary dynamical system.

In practice, most efforts to explain variable tFUS responses have focused on the stimulation itself by mapping dose-response curves across acoustic parameters [36, 56]. Less attention has been given to the choice of outcome measure. The dependent variable is typically a measure of overall activity, such as population firing rate or fluorescence amplitude. These readouts quantify how much activity changed, but are largely agnostic to the underlying temporal organization. In a dynamical system, an amplitude increase can arise from different regimes: a tonic elevation, a transient burst, or a shift in excitatory-inhibitory balance. Moreover, dynamical reorganization and amplitude changes need not co-occur: a perturbation can alter temporal organization without producing a resolvable change in mean amplitude, or vice versa.

This limitation motivates a complementary viewpoint from nonequilibrium statistical physics. The brain is a paradigmatic far-from-equilibrium system that continuously expends energy to maintain organized activity patterns. tFUS is itself an energetic perturbation: it deposits acoustic energy into dissipative neural tissue. Understanding neuromodulation is thus partly a question of how a system that is already far from thermodynamic equilibrium processes an additional, exogenous energy flux. The field of stochastic thermodynamics [16, 47] provides the framework to address this question. Central to this framework are fluctuation theorems [11, 22] that relate nonequilibrium work to equilibrium free-energy differences, and estimators of entropy production rate (EPR) that quantify broken detailed balance and have revealed nonequilibrium dynamics in living systems [3, 20]. In systems weakly coupled to an environment, stochastic thermodynamics relates fluctuations, time-irreversibility, and entropy production, connecting observed trajectories to dissipation and the thermodynamic arrow of time [13, 42, 46]. Recent work has shown that temporal irre-versibility estimated from neural time series captures mesoscopic nonequilibrium structure in the brain, with irreversibility varying across brain states, arousal levels, and pharmacological conditions [12, 31, 32, 45]. If sonication changes the nonequilibrium character of neural dynamics (i.e., how irreversibly the system evolves in time) then stochastic thermodynamics may afford a new understanding of ultrasonic neuromodulation.

Here we employ tools from stochastic thermodynamics to conduct a novel analysis of an existing dataset from Murphy et al. [36], which combined fiber photometry of the behaving rodent thalamus with tFUS at varying intensities (Figure 1A,B). We estimated an ordinal surrogate of the entropy production rate (EPR) before, during, and after stimulation (Figure 1C,D) to address the following questions: (i) Does tFUS modulate neural irreversibility either during or immediately after sonication? (ii) Does EPR contain information beyond that of calcium amplitude? (iii) Does prestimulation irreversibility predict the subsequent response to stimulation? We present evidence that all three answers are affirmative and discuss their implications for the mechanistic understand-ing of ultrasonic neuromodulation.

**Figure 1:**
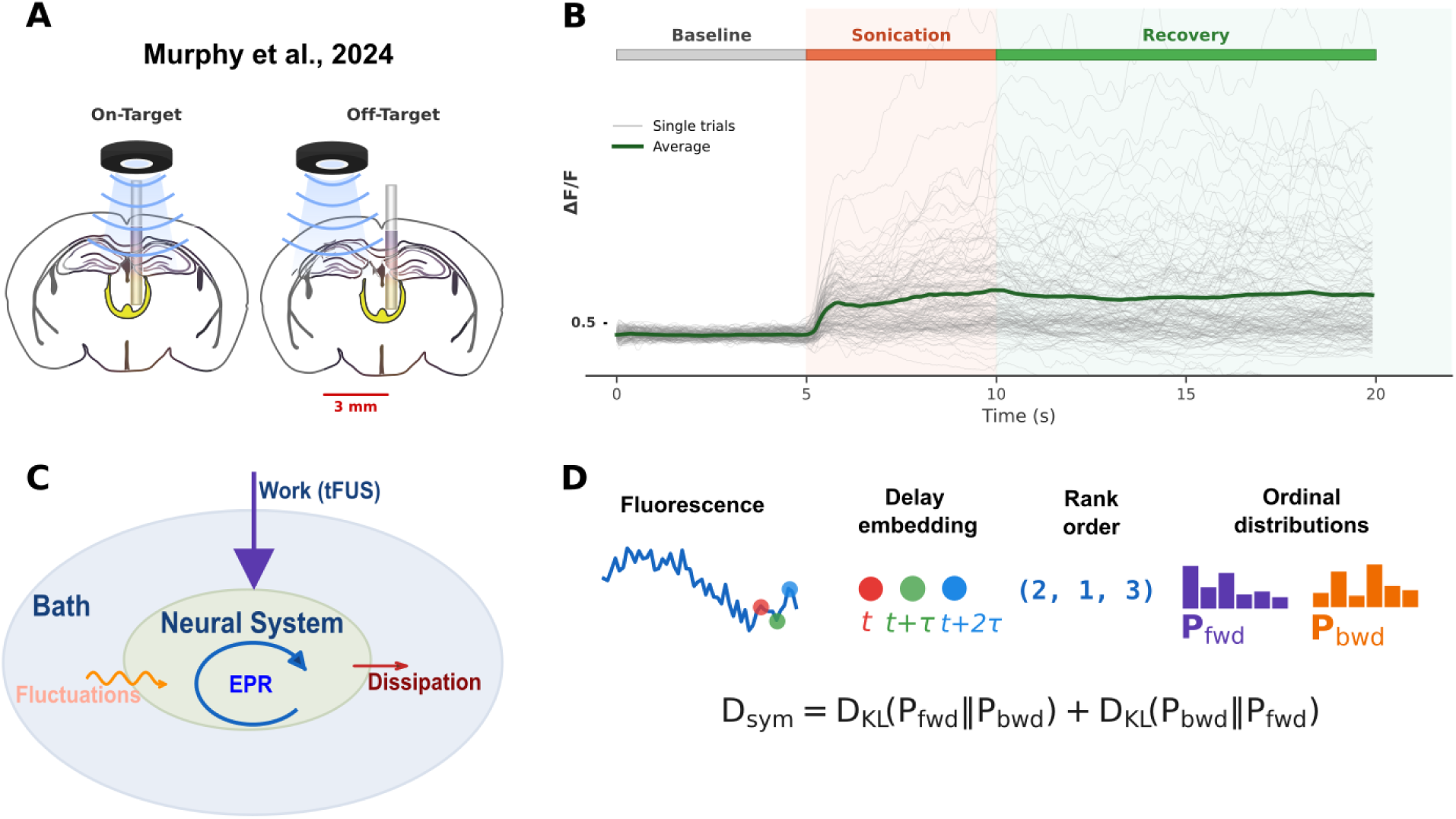
A stochastic thermodynamics framework for investigating neural responses to focused ultrasound. **(A)** Experimental setup from Murphy et al. [36]. A scalp transducer (550 kHz) delivered tFUS to the central medial thalamus (CMT) of freely moving GCaMP-expressing mice while neural activity was recorded via fiber photometry. Stimulation was applied on-target (directly above the CMT) and off-target (3 mm lateral offset). **(B)** Fluorescence traces. Each trial consisted of a 5 s baseline, 5 s sonication, and recovery period (first 10 s analysed). Gray traces show individual trials; the dark green trace shows the trial average. **(C)** Stochastic thermodynamics framework. tFUS is modeled as an external work input to a neural system that is already far from thermodynamic equilibrium and weakly coupled to an effective bath. The resulting stochastic dynamics reflect bath-driven fluctuations, internal entropy production, and heat dissipation back to the environment. **(D)** Ordinal EPR estimation pipeline. The calcium signal is delay-embedded, and each embedding vector is mapped to its ordinal pattern. The symmetrised Kullback–Leibler divergence between the forward and backward ordinal distributions yields *D*_sym_, an estimator of the EPR.

## 2 Results

### 2.1 Calcium amplitude and irreversibility show distinct dose-response pat-terns

We analyzed fiber photometry recordings from the central medial thalamus (CMT) [36]. Each ana-lyzed trial included a 5 second baseline, 5 seconds of stimulation, and the 10 seconds immediately after tFUS (“recovery”; Figure 1B). Pulsed tFUS (pulse repetition frequency PRF = 2.5 Hz; carrier frequency 550 kHz) was applied either to the recorded region (“on-target”) or to the area 3 mm laterally adjacent to the CMT (“off-target”; Figure 1A). The acoustic intensity spanned six values ranging from 0.3 to 7.4 W/cm^2^. We estimated an ordinal surrogate of the EPR (Figure 1C-D) after pooling trials within each animal-condition (see “Pooled-trial EPR and calcium amplitude measures” in *Methods*). To evaluate dose-response for both EPR and GCaMP amplitude, we employed mixed-effects models with a common specification: the baseline value was included as a covariate, and the predictors of each model included the condition (on-target versus off-target) and acoustic intensity (both linear and quadratic terms; see Eq. 4).

Calcium amplitude increased monotonically with dose in the on-target condition during both stimulation and recovery, while off-target fluorescence remained low across the intensity range (Fig. 2C,D,F). On-target stimulation produced significantly larger calcium responses than off-target in both windows (stimulation: *p <* 0.001; recovery: *p <* 0.001). On-target-specific dose structure was significant during stimulation, with both the linear (*β*_on×*I*_ = +7.57, *p* = 0.006) and quadratic (*β*_on×*I*2_ = −48.3, *p* = 0.010) interaction terms reaching significance, consistent with a monotonic increase that saturates at higher intensities. During recovery, calcium amplitude showed only a linear dependence on intensity (*β*_on×*I*_ = +16.34, *p <* 0.001; quadratic *p* = 0.746).

**Figure 2:**
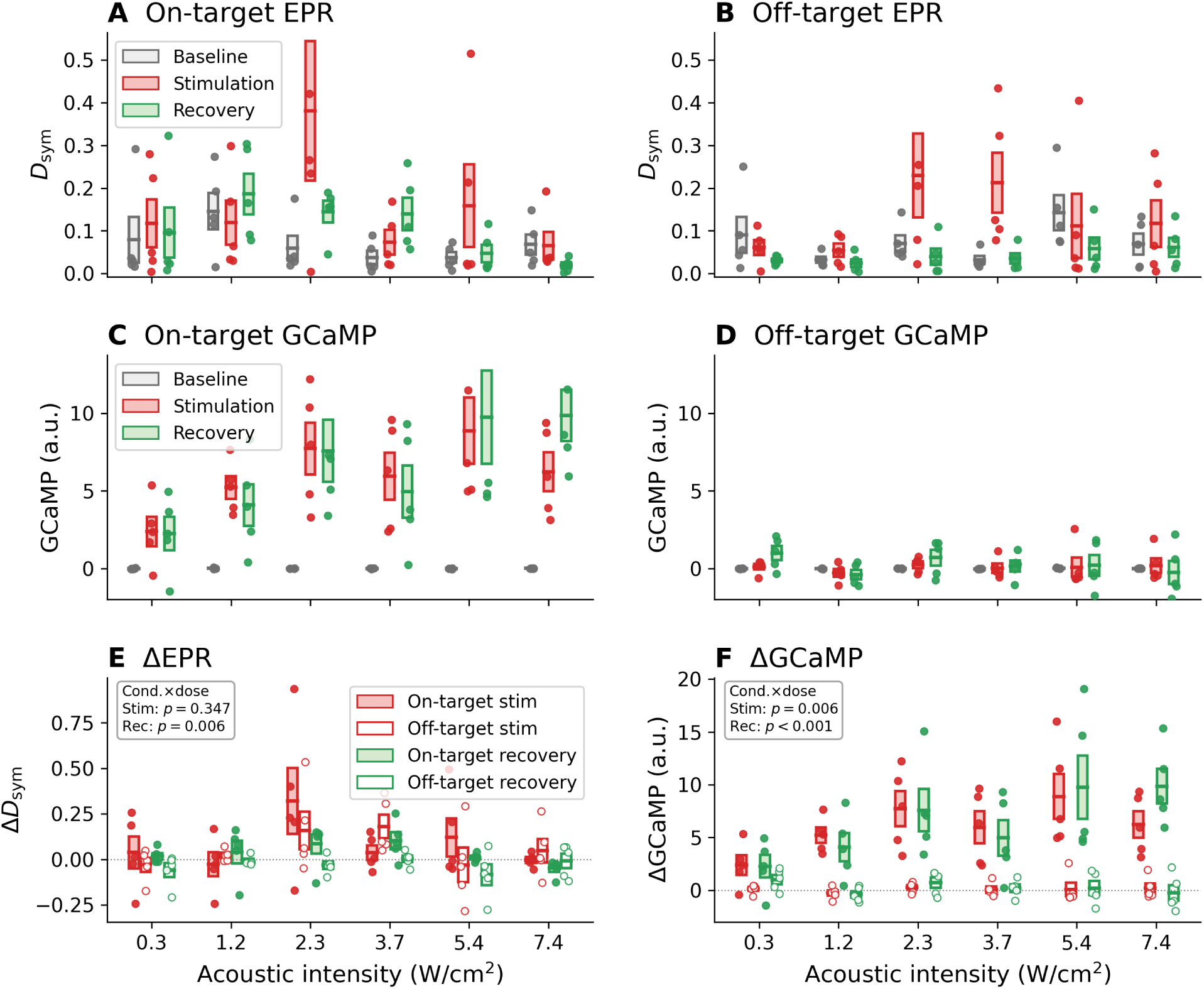
Distinct dose-response of temporal irreversibility and calcium amplitude. We mea-sured the change in both EPR and GCaMP during and immediately after 2.5 Hz sonication of the central medial thalamus (CMT). Stimulation was delivered on-target and with a 3 mm off-set (off-target) between tFUS and GCaMP acquisition. **(A,B)** EPR (*D*_sym_) in the on-target (A) and off-target (B) conditions during baseline (grey), stimulation (red), and recovery (green). The largest deviations from baseline are found at moderate intensities. Interestingly, the non-monotonic dose-response is also apparent (but muted) during off-target stimulation, despite the absence of a corresponding calcium response. **(C,D)** GCaMP amplitude in the on-target (C) and off-target (D) conditions. Unlike EPR, GCaMP shows a largely monotonic dose-response, with no response during off-target sonication. **(E)** Baseline-subtracted EPR (Δ*D*_sym_), shown for on-target/off-target stimulation and recovery. Contrast between on-and off-target is largest in the recovery period. **(F)** Baseline-subtracted GCaMP (ΔGCaMP). The contrast between on-and off-target increases with intensity. Boxes show the mean ±1 SEM across animals; points denote individual animals.

Temporal irreversibility showed a qualitatively different dose dependence. In the on-target condition (Fig. 2A,E), both the stimulation and recovery windows showed the largest changes from baseline at moderate intensities (2.3–3.7 W/cm^2^). Interestingly, the off-target condition (Fig. 2B,E) also showed a non-monotonic irreversibility response (albeit milder) despite the absence of a strong fluorescence signal, hinting that sonication can alter dynamics in the absence of an amplitude change.

During stimulation, we did not resolve an on-target main effect for EPR (*p* = 0.690), nor did we find significant condition-intensity interactions (*p* = 0.216 and *p* = 0.872 for the linear and quadratic interactions, respectively). The absence of a significant on-target contrast during stimulation is consistent with the observation that *off-target* sonication also elevated EPR in this window (Fig. 2B,E). In recovery, by contrast, on-target EPR was significantly higher than off-target (*p <* 0.001), and both on-target-specific dose interaction terms were significant (*β*_on×*I*_ = −0.265, *p* = 0.002; *β*_on×*I*2_ = −1.550, *p* = 0.006). Note that a negative regression coefficient on the quadratic dose term is consistent with an inverted-U response. Overall, the effect of on-target stimulation on EPR emerges immediately after sonication. Table 1 summarises the on-target-specific terms for both readouts and both windows.

**Table 1:** Dose-response relationships for calcium amplitude and EPR. Each readout was modelled with a mixed-effects model following Eq. 4. Only the on-target main effect and its dose interactions are shown; these terms capture on-target-specific effects of acoustic intensity on the corresponding outcome measure. ^∗^ *p <* 0.05; ^∗∗^ *p <* 0.01; ^∗∗∗^ *p <* 0.001.

| Window | Term | EPR ($D_{\text{sym}}$) | | | | GCaMP (fluorescence) | | | |
| --- | --- | --- | --- | --- | --- | --- | --- | --- | --- |
| | | $\hat{\beta}$ | SE | $z$ | $p$ | $\hat{\beta}$ | SE | $z$ | $p$ |
| Stim. | on-target | +0.027 | 0.069 | 0.40 | .690 | +7.63 | 0.86 | 8.87 | <.001*** |
| | on-target $\times I_c$ | -0.329 | 0.266 | -1.24 | .216 | +7.74 | 3.15 | 2.46 | .014* |
| | on-target $\times I_c^2$ | -0.286 | 1.774 | -0.16 | .872 | -47.7 | 21.7 | -2.20 | .028* |
| Rec. | on-target | +0.108 | 0.022 | 4.80 | <.001*** | +6.78 | 1.09 | 6.24 | <.001*** |
| | on-target $\times I_c$ | -0.265 | 0.085 | -3.12 | .002** | +16.5 | 3.96 | 4.17 | <.001*** |
| | on-target $\times I_c^2$ | -1.550 | 0.569 | -2.72 | .006** | -5.15 | 27.3 | -0.19 | .850 |

### EPR dose-response is consistent across animals

The inverted-U EPR dose-response pattern during recovery was clearly visible in 4 of 5 animals, with the peak change from baseline (Δ*D*_sym_) occurring between 1.2 and 3.7 W/cm^2^ (Fig. S1). The pattern was not observed with off-target stimulation (Fig. S2). Thus, the quadratic group-level effect was not driven by a single strong “responder” and was specific to sonication of the CMT.

### No significant GCaMP response at 20 Hz PRF

Murphy et al. [36] reported an offline inhibitory effect when stimulating the CMT with a higher PRF of 20 Hz. We were not able to resolve a main effect of on-target stimulation on calcium amplitude during stimulation (*p* = 0.676) or recovery (*p* = 0.821), finding instead that the change in GCaMP remained positive (Fig. S3). We discuss potential reasons for this discrepancy in the Discussion. The corresponding analysis of EPR did not resolve a significant main effect of on-target sonication during stimulation (*p* = 0.940), although a nominally significant EPR *reduction* was present during recovery (*p* = 0.044).

### 2.2 Irreversibility cannot be explained by calcium amplitude alone

Figure 3 depicts the change from baseline for both EPR and GCaMP, shown separately during (panel A) and after (panel B) stimulation. The sharp falloff of irreversiblity at higher acoustic intensities, especially during recovery, suggests that EPR is capturing a distinct aspect of CMT activity. We thus measured the correlation between the two assays of neural response – Δ*D*_sym_ and ΔGCaMP – during and immediately after on-target tFUS (Fig. 3C-D, respectively).

**Figure 3:**
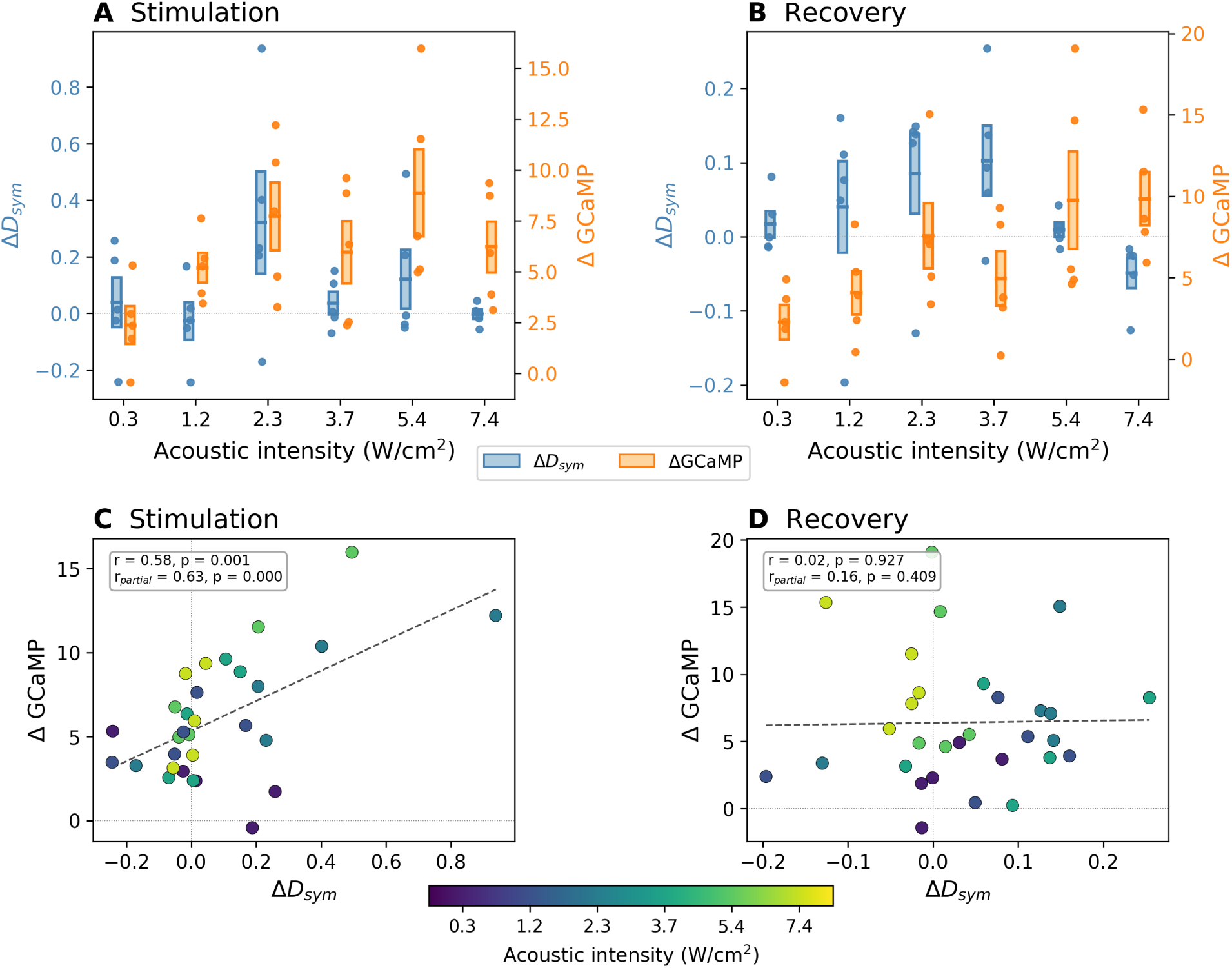
Dissociation between irreversibility and calcium amplitude immediately after son-ication. **(A)** Dual-axis overlay of on-target Δ*D*_sym_ (blue, left axis) and ΔGCaMP (orange, right axis) during stimulation. **(B)** Same as (A) but now shown for the recovery period. The change in EPR peaks at moderate intensities while calcium amplitude is largely monotonic with dose. Error bars: ±1 SEM. **(C)** Scatter of Δ*D*_sym_ versus ΔGCaMP during stimulation (each marker corresponds to one animal×intensity combination, *n* = 30). Raw and partial correlations are both positive (*r* = 0.58, *p <* 0.001; *r*_partial_ = 0.63, *p <* 0.001). **(D)** During recovery, the correlation vanishes (*r* = 0.02, *p* = 0.927; *r*_partial_ = 0.16, *p* = 0.409), indicating that post-sonication EPR is decoupled from fluorescence amplitude despite both measures being significantly elevated relative to baseline.

During stimulation, the changes from baseline were positively correlated (*r* = 0.58, *p <* 0.001), even after conditioning the correlation on intensity (*r*_partial_ = 0.63, *p <* 0.001). During recovery, however, the correlation vanished (*r* = 0.02, *p* = 0.927; *r*_partial_ = 0.16, *p* = 0.409), despite the fact that both EPR and GCaMP are significantly different from baseline levels during the recovery period (Table 1).

To test whether recovery EPR (Fig. 4B) carries dose information that calcium amplitude cannot account for, we fit a set of mixed models (see “EPR beyond calcium amplitude” in *Methods*) where the base (reduced) model captures the dependence of EPR on its baseline value *and* the observed change in GCaMP (Fig. 4A). Adding linear and quadratic dose terms significantly increased model fit (on-target stimulation trials only; *χ*^2^(2) = 13.55, *p* = 0.001), indicating the presence of dose structure beyond calcium amplitude. When pooling on-target and off-target trials and adding dose interaction terms, model fit was further improved (*χ*^2^(3) = 27.23, *p <* 0.001). Importantly, the ΔGCaMP covariate was not significant (*β* = −0.006, *p* = 0.642) in the full model, indicating that the level of irreversibility during recovery is not associated with the change in calcium amplitude. We calculated a residual EPR by regressing out its baseline and the associated change in GCaMP amplitude. The residualized EPR is shown as a function of dose in Fig. 4C. In the on-target condition, the residual is positive from 0.3 to 3.7 W/cm^2^ but negative at 5.4 and 7.4 W/cm^2^. In other words, high-dose sonication produced less irreversibility than expected from its calcium-amplitude response.

**Figure 4:**
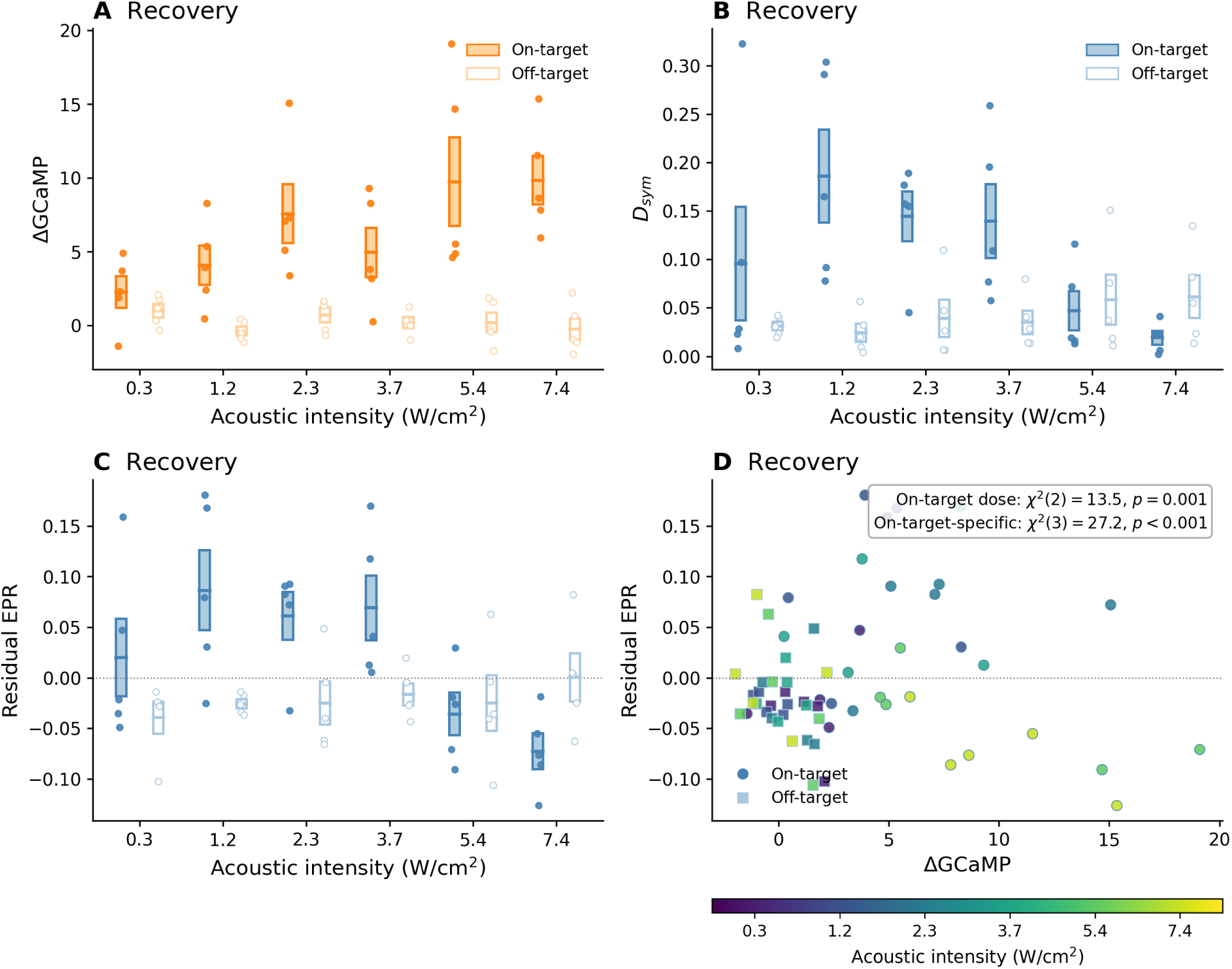
EPR cannot be explained by calcium amplitude. To identify whether changes in EPR can be accounted for by concomitant changes in calcium amplitude, we analyzed the relationship between the change in GCaMP (recovery versus baseline), baseline irreversibility, and the irre-versibility measured during recovery. **(A)** The change in GCaMP after sonication (ΔGCaMP) rises strongly with dose after on-target tFUS. **(B)** The EPR after sonication (*D*_sym_) peaks at interme-diate intensity and falls off at higher doses. **(C)** The residual EPR after regressing out ΔGCaMP and baseline EPR: the residual is positive at low-to-moderate on-target doses and negative at the highest on-target doses. Error bars: ±1 SEM across animals. **(D)** Residual EPR versus ΔGCaMP, coloured by intensity. We fit a set of mixed models to determine whether the residual EPR has a dose-dependence, and if so, whether the dose-dependence is specific to on-target sonication. Within the on-target condition, dose remained significant after accounting for baseline EPR and ΔGCaMP (*χ*^2^(2) = 13.55, *p* = 0.001). Allowing dose terms to differ between on-target and off-target further improved fit (*χ*^2^(3) = 27.23, *p <* 0.001).

We repeated the hierarchical mixed model analysis for the EPR observed during the *stimulation* period (Fig. S4), and again found that dose improved fit beyond baseline EPR the change in GCaMP (on-target only; *χ*^2^(2) = 7.28, *p* = 0.026). As in recovery, the model fit was significantly improved with the addition of on-target versus off-target terms (*χ*^2^(3) = 15.27, *p* = 0.002). However, unlike recovery, the ΔGCaMP covariate was statistically significant (*β* = +0.181, *p <* 0.001). In summary, after accounting for baseline value and the change in calcium amplitude, EPR still retains significant dose structure, indicating that sonication reshapes recovery dynamics beyond what amplitude alone would predict. Complete likelihood-ratio test statistics are provided in Table S1.

### 2.3 Prestimulation irreversibility predicts stimulation responsiveness

We asked whether the prestimulation level of temporal irreversibility – a single-trial marker of neural state – predicted the magnitude or direction of the response to tFUS. Both EPR and GCaMP responses were computed at the single-trial level for baseline, stimulation, and recovery windows across all six acoustic intensities (*n* = 408 trials for 2.5 Hz PRF).

#### Baseline state modulates stimulation EPR

To test whether baseline irreversibility influences the subsequent change in EPR during or after sonication, we partitioned trials into terciles based on their prestimulation EPR. Trials were pooled across intensities and condition such that we considered both on-target and off-target stimulation of the CMT. We then fit a mixed-effects model with baseline-EPR tercile, condition, and their interaction as fixed effects, with a random intercept for animal (see “Baseline-state mixed-effects models” in *Methods*). During stimulation, we found a significant baseline-state × condition interaction (*χ*^2^(2) = 11.49, *p* = 0.003), indicating that the relationship between baseline EPR and the subsequent change differs significantly when stimulating the recorded region (Figure 5A). More specifically, on-target stimulation generated a larger increase in EPR when prestimulation irreversibility was *lower*. In recovery, the corresponding interaction was absent (*χ*^2^(2) = 1.05, *p* = 0.592; Figure 5B), indicating that the baseline-state dependence of EPR is concentrated in the stimulation period rather than persisting into recovery.

**Figure 5:**
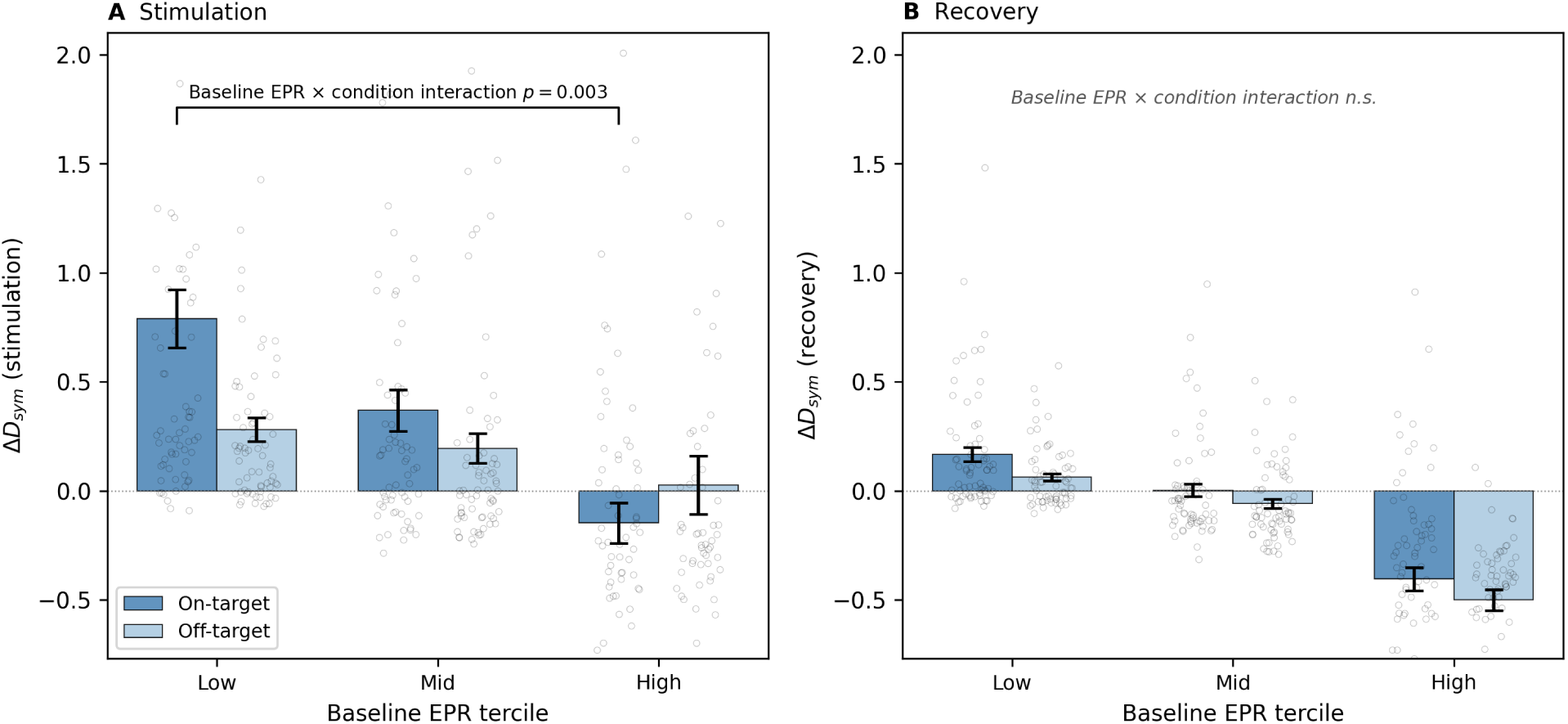
Baseline irreversibility modulates EPR response during but not after stimulation. Vertical axes show the change in irreversibility for on-target (dark blue) and off-target (light blue) trials pooled across all six acoustic intensities and stratified by baseline EPR tercile. **(A)** During stimulation, the change in EPR is significantly modulated by baseline irreversibility (baseline-EPR × condition interaction: *p* = 0.003). Trials with lower prestimulation EPR respond more strongly to tFUS in a target-specific manner. **(B)** In contrast, the change in EPR measured dur-ing recovery is not significantly modulated by baseline irreversibility (*p* = 0.592). Error bars: ±1 SEM.

#### Baseline EPR modulates the change in GCaMP

We also investigated whether prestimulation irreversibility predicts the change in *GCaMP* during and after stimulation (see “Baseline-state mixed-effects models” in *Methods*). During stimulation, we found a significant interaction between baseline EPR and condition (*χ*^2^(2) = 9.19, *p* = 0.010), indicating that the baseline EPR dependence of the calcium response differed between on-target and off-target tFUS. In recovery, the corresponding interaction was not significant (*p* = 0.113). Baseline EPR also contributed to overall GCaMP variability beyond condition and intensity, reaching significance in recovery (*χ*^2^(2) = 6.45, *p* = 0.040) and trending similarly during stimulation (*χ*^2^(2) = 5.56, *p* = 0.062).

Figure 6C,D shows the corresponding adjusted means: on-target responses are largest in the low-baseline tercile and reduced in the high-baseline tercile, whereas off-target responses remain near zero across strata.

**Figure 6:**
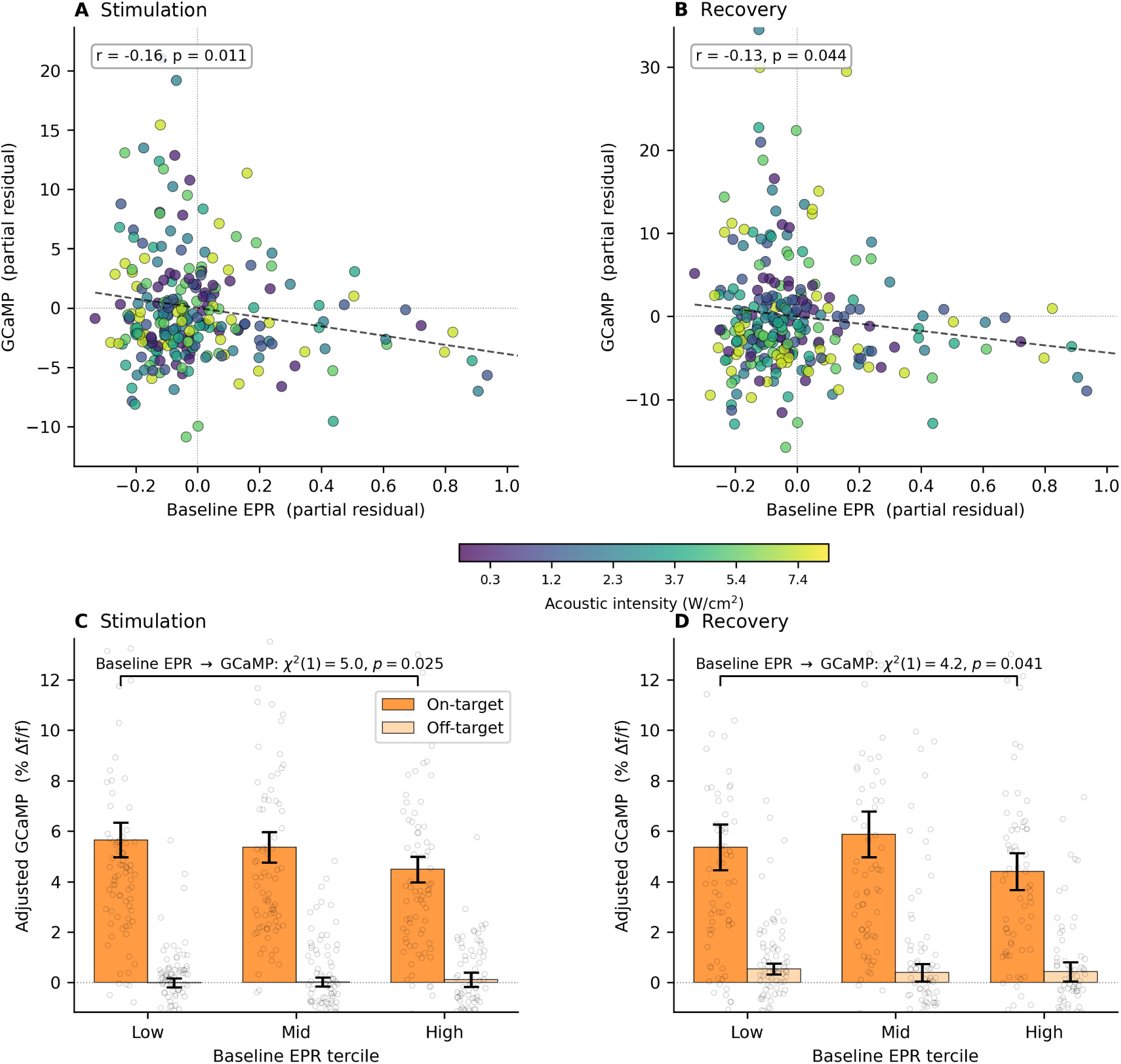
Baseline EPR predicts GCaMP response beyond baseline GCaMP and intensity. (A,B) Added-variable (partial regression) plots for stimulation (A) and recovery (B), shown for on-target trials only. Each point is a single trial coloured by acoustic intensity; axes show residu-als after removing baseline GCaMP, intensity, and animal random effects from both baseline EPR (x-axis) and GCaMP (y-axis). The negative slope indicates that higher prestimulation irreversibil-ity predicts weaker calcium responses. **(C,D)** Mean adjusted GCaMP (residualized for baseline GCaMP and intensity) by baseline-EPR tercile during stimulation (C) and recovery (D) for on-target (dark orange) and off-target (light orange) conditions. After removing the effects of baseline amplitude and dose, on-target responses still decrease with baseline EPR (likelihood-ratio test for adding baseline EPR: stimulation *χ*^2^(1) = 5.04, *p* = 0.025; recovery *χ*^2^(1) = 4.17, *p* = 0.041). Error bars: ±1 SEM.

To quantify the incremental predictive value of baseline EPR, we fit trial-level on-target-only mixed models in which GCaMP was predicted from baseline GCaMP, acoustic intensity (linear and quadratic), and a random animal intercept (see “Incremental prediction of GCaMP by baseline EPR” in *Methods*). Adding baseline EPR to that model significantly improved fit during stimulation (*χ*^2^(1) = 5.04, *p* = 0.025; *β* = −2.78, Δ*R*^2^ = 0.020) and during recovery (*χ*^2^(1) = 4.17, *p* = 0.041; *β* = −3.54, Δ*R*^2^ = 0.017). In both windows, the sign was negative: higher prestimulation irreversibility predicted a weaker calcium response after controlling for baseline calcium and dose. Figure 6 shows this relationship via added-variable (partial regression) plots and adjusted tercile means.

Thus, baseline EPR predicts overall response magnitude, with lower-baseline trials showing larger calcium responses on average – a relationship that holds after adjusting for baseline GCaMP and acoustic intensity.

### 2.4 Sensitivity to ordinal parameters

Because ordinal EPR depends on the embedding dimension *m* and delay *τ*, we tested the influence of parameter choices on our findings. For the main analysis presented above, we selected *m* = 3 to minimize the dimensionality of the forward and reverse probabilities, which increase with *m*!. The value for the delay *τ* = 8 was selected from a split-half reliability sweep of baseline irreversibility across *τ* = {2, 4, 6, 8}.

The core findings were preserved with (*m, τ*) = (3, 6). During recovery, EPR dose-response showed a significantly negative quadratic dose relationship (*β_I_*_2_ = −1.90, *p* = 0.032). The EPR during recovery was shown to contain information beyond that of calcium amplitude (on-target-specific dose: *χ*^2^(3) = 17.50, *p* = 5.6 × 10^−4^; on-target-only dose: *χ*^2^(2) = 8.75, *p* = 0.013; ΔGCaMP covariate remained null: *p* = 0.992). Baseline EPR significantly influenced the subsequent change in both EPR (baseline-EPR × condition interaction: *χ*^2^(2) = 7.00, *p* = 0.030) and GCaMP amplitude (*χ*^2^(2) = 6.70, *p* = 0.035) during stimulation.

The main claims were also largely preserved with a higher-dimensional embedding, namely (*m, τ*) = (4, 8). The EPR during recovery exhibited a significant quadratic dose dependence (*β_I_*_2_ = −2.12, *p* = 0.015), and both on-target-specific recovery dose terms remained significant (*β*_on×*I*_ = −0.39, *p* = 0.002; *β*_on×*I*2_ = −2.12, *p* = 0.015). Moreover, calcium amplitude could not explain the EPR measured during recovery (on-target-specific dose: *χ*^2^(3) = 24.59, *p* = 1.9 × 10^−5^; on-target-only dose: *χ*^2^(2) = 11.07, *p* = 0.0040; ΔGCaMP covariate again remained null: *p* = 0.911). Baseline EPR predicted the subsequent change in irreversibility during sonication in a target-specific manner (*χ*^2^(2) = 12.59, *p* = 0.0018). Baseline irreversibility tended to also predict the associated change in calcium amplitude during sonication, although the effect did not reach significance (*p* = 0.105).

## 3 Discussion

By employing tools from stochastic thermodynamics, this study found that low-intensity focused ultrasound perturbs brain activity in a manner that is not captured by conventional amplitude-based readouts. The GCaMP signal recorded via fiber photometry reflects aggregate calcium dynamics across the illuminated neural population, encompassing both somatic and neuropil compartments [9, 26]. When stimulating the CMT with tFUS, the amplitude of the GCaMP signal increased monotonically with the intensity of the acoustic stimulus [36] (Fig. 2). By contrast, the EPR showed a non-monotonic dose-response that peaked at moderate intensity (Figure 2A,B,E), a pattern that was visible in both on-target and off-target conditions.

### An inverted-U dose–response for temporal irreversibility

The inverted-U pattern of EPR is reminiscent of nonlinear dose-response relationships that have been documented across several forms of brain stimulation and, more broadly, across biology. The classical Yerkes–Dodson law posits that behavioural performance peaks at intermediate levels of arousal or stimulus intensity [52], and the hormesis literature documents biphasic dose-response curves in which low doses stimulate and high doses inhibit across a wide range of biological measures [8]. In transcranial direct-current stimulation, moderate current densities produce the strongest effects on cortical excitability and cognition, whereas higher intensities can reverse or abolish these effects [2, 15]. Deep brain stimulation of the central thalamus likewise enhances working memory at low currents and impairs it at higher currents [33]. The concept of stochastic resonance – where an intermediate level of noise optimally enhances signal detection in a nonlinear system – provides a formal framework for understanding such inverted-U phenomena [19, 35]. The findings here point to the possibility that the brain’s response to exogenous energy input is fundamentally nonlinear, with an intermediate regime in which stimulation most effectively engages endogenous dynamics.

For ultrasound, one interpretation is that moderate acoustic doses perturb the system enough to displace it from its resting attractor without overwhelming the endogenous dynamics that generate temporal irreversibility. At higher intensities, the exogenous drive may dominate the local dynamics – producing large calcium transients – while constraining the system to a more stereotyped, less dissipative trajectory. In the language of nonequilibrium physics, moderate sonication may optimally couple to the system’s internal degrees of freedom, generating the greatest departure from time-reversal symmetry. By contrast, strong sonication may effectively “entrain” the population [21, 41], increasing the amplitude of the response while reducing its dynamical complexity. Although speculative, this framing makes the testable prediction that the peak of the EPR dose-response should shift with parameters that alter the balance between exogenous drive and endogenous dynamics, such as pulse repetition frequency or duty cycle.

### The post-sonication dissociation between irreversibility and amplitude

It is notable that the dissociation between GCaMP and EPR is most pronounced immediately *after* sonication (Figure 3B,D; Figure 4). During this period, the system is reorganising after perturbation, and both measures remain elevated above their respective baselines. The fact that the two measures are uncorrelated during this window (Figure 3D) indicates that amplitude and irreversibility reflect distinct aspects of the region’s relaxation. In other words, the path by which the system relaxes after perturbation carries information that is invisible to the amplitude readout. Moderate-intensity sonication may push the system into a region of state space from which the return to baseline involves the greatest thermodynamic cost – the largest entropy production — whereas high-intensity stimulation, despite producing larger fluorescence changes, may push the system along a more direct, less dissipative relaxation path.

### Off-target stimulation elevates irreversibility without a calcium response

An interesting as-pect of the present findings is that stimulating a region 3 mm adjacent to the site of the recording fiber produced a clear inverted-U EPR elevation during stimulation that was visible in multiple animals (Figs. 2B, S2). In the mouse brain, a 3 mm offset from the CMT places the acoustic focus well outside of the thalamus proper [40], yet the two sites remain within reach of common thalamocortical and corticothalamic projection systems. It is therefore plausible that off-target sonication perturbs polysynaptically connected circuits whose altered dynamics propagate to the recorded CMT population. If these network-level perturbations produce changes in temporal structure without substantially increasing the aggregate calcium flux, they would appear in the irreversibility measure but not in the bulk fluorescence signal. This interpretation is consistent with the broader notion that EPR is sensitive to changes in the temporal organization of activity that need not manifest as amplitude changes. Note that for conventional neuromodulation studies, off-target stim-ulation serves as an experimental control. Here, it represents an informative finding in of itself, suggesting that sonication can alter the nonequilibrium character of neural dynamics at a distance, and that EPR may be sensitive to low-amplitude perturbations.

### Baseline irreversibility as a susceptibility marker

Another aspect of our findings suggests that EPR serves as a proxy for neural “susceptibility”: trials with lower baseline irreversibility showed systematically higher responses during stimulation, measured with both GCaMP and EPR itself (Figs. 5, 6). This is conceptually aligned with findings in other stimulation modalities showing that prestimulus brain state governs the response to transcranial stimulation. In TMS, for example, cortical excitability tracks the instantaneous phase of endogenous oscillations [44], and state-dependent protocols can selectively induce plasticity that is absent when stimulation is delivered irrespective of brain state [4, 58]. The EPR effect – lower baseline *D*_sym_ predicting larger Δ*D*_sym_ – could in principle reflect regression to the mean, since a high-baseline trial will mechanically tend toward a negative change score. However, our data argue against such an artifactual explanation. The relationship between baseline irreversibility and the subsequent change was significantly modulated by whether tFUS was applied on-or off-target (Figure 5A). Moreover, baseline EPR predicted the GCaMP *amplitude* response even after controlling for baseline GCaMP and acoustic intensity (Figure 6). This cross-modal prediction (i.e., prestimulation irreversibility anticipates the magnitude of the calcium-amplitude response) is evidence that baseline EPR captures a genuine axis of stimulation susceptibility.

### No effects when stimulating at 20 Hz

Murphy et al. [36] reported that 20 Hz PRF stimulation of the CMT produced an offline, inhibitory calcium response. While our analysis of 2.5 Hz stim-ulation was highly congruent with that of Murphy et al. [36], we did not find significant changes in either calcium amplitude or EPR at a PRF of 20 Hz (Supplementary Fig. S3). This discrepancy may stem from differences in the preprocessing that were employed in the respective studies, particularly the approaches used to correct for motion artifacts. The finding that 20 Hz stimulation did not modulate irreversibility is consistent with the interpretation that the 20 Hz protocol elicited an overall weaker neuromodulatory effect compared to 2.5 Hz.

### Limitations: thermodynamic interpretation

Our employment of an ordinal surrogate of EPR (*D*_sym_; see Eq. 2) draws on the formal connection between time-reversal asymmetry and dissipation in nonequilibrium statistical physics. The foundational result – that the Kullback–Leibler divergence between forward and reverse trajectory probabilities equals the mean entropy produc-tion – was established for Markovian systems coupled to a thermal bath [11, 22, 47] and extended to discrete-state systems [16, 42, 43]. Practical estimators that detect broken detailed balance from time-series data have extended the applicability of this framework [34, 46]. Broken detailed balance has been identified at mesoscopic scales in active biological matter [3, 20]. Applications to neural data have shown that human brain dynamics violate detailed balance at large spatial scales, with the degree of irreversibility tracking arousal state and cognitive load [12, 31, 32, 45].

However, neural tissue violates the assumptions of these theorems in several ways. The GCaMP signal reflects calcium dynamics filtered through an indicator with slow kinetics. The system has multiple chemical reservoirs, and the ordinal embedding captures only a low-dimensional projection of the underlying state space. The symmetrised KL divergence used here is therefore best understood as a nonparametric measure of time-arrow asymmetry in the observed signal: a quantity that is related to, but not identical with, the thermodynamic EPR of the underlying biophysical system. Nevertheless, our results indicate that this time-asymmetry measure carries information about the neural response to sonication that is inaccessible to conventional amplitude readouts.

### Limitations: EPR estimation

EPR estimation from noisy biological time series is intrinsically challenging. Any KL-based irreversibility estimator depends on the embedding parameters and on finite-sample bias, especially in short windows. We employed an ordinal-based method because it is nonparametric, insensitive to monotone rescalings of the signal, and naturally suited to short segments [1, 53]. Alternative approaches to estimating entropy production from time series include visibility-graph irreversibility [24], neural-network classifiers trained on the arrow of time [46], model-based estimators that infer dissipation from partially observed Langevin dynamics [34], and current-based methods that quantify dissipation from fluctuating probability currents [28]; a recent review provides a systematic comparison [54]. Analytical treatments of EPR in neural-field models have also begun to connect irreversibility to network parameters [30].

### Broader implications

Nonequilibrium statistical physics offers a language for asking how an exogenous stimulus does work on a living system that is already far from thermodynamic equilibrium. In this framework, sonication does not merely increase or decrease neural activity; it can also reshape the path which the system takes during and after perturbation. The present results suggest that moderate doses may drive the system into a regime with greater temporal irreversibility than would be expected from fluorescence amplitude alone, whereas higher doses can produce large calcium increases with less irreversibility. Future work could test this more directly by combining time-asymmetry markers with electrophysiology and metabolic imaging to establish whether irreversibility changes co-occur with shifts in biophysical dissipation, and by testing closed-loop stimulation designs that target different baseline states [29, 57]. More generally, our findings suggest that stochastic thermodynamics serves as a valuable complement to conventional amplitude-based readouts in neuromodulation, as it captures the interaction between exogenous energy and the brain’s ongoing nonequilibrium dynamics.

## 4 Methods

### 4.1 Dataset

We analysed fiber photometry recordings from the publicly available dataset of Murphy et al. [36], which investigated the neuromodulatory effects of transcranial focused ultrasound (tFUS) in the mouse deep brain. Data was accessed via https://dataverse.harvard.edu/dataset. xhtml?persistentId=doi:10.7910/DVN/PCWRAD. In brief, adult C57BL/6J mice were injected with adeno-associated viral vectors encoding the genetically encoded calcium indicator GCaMP6s [9] under the CaMKII promoter and implanted with 200 *µ*m-diameter optical fibers targeting the thalamic central medial nucleus (CMT). Ultrasound was delivered through the intact skull using a wearable, miniaturised ring transducer operating at a fundamental frequency of 550 kHz with a lateral focal spot of ∼2.3 mm (full-width at half-maximum). The transducer was chronically mounted on a 3D-printed head frame, enabling repeated stimulation sessions in freely moving animals.

Dual-wavelength fiber photometry was performed using interleaved 470 nm (calcium-dependent GCaMP6s excitation) and 405 nm (calcium-independent isosbestic) illumination, providing a noise reference for correcting motion and hemodynamic artifacts. Data was sampled at 32 Hz. Each recording session comprised multiple trials at six output levels corresponding to reported spatial-peak pulse-average acoustic intensities of 0.3, 1.2, 2.3, 3.7, 5.4, and 7.4 W/cm^2^. A trial consisted of a 5-second prestimulation baseline, a stimulation epoch (5 s when stimulating at 2.5 Hz PRF; 40 s at 20 Hz PRF), and a post-stimulation recovery period such that the total length of each trial was 180 s. For our analysis, we retained only the first 10 second of the recovery period.

Stimulation was conducted on-target (sonication directed at the recorded CMT region) and separately off-target (sonication of the area 3 mm laterally adjacent to the CMT; Fig. 1A). For each configuration, two pulse repetition frequencies (PRFs) were tested: 2.5 Hz and 20 Hz. Murphy et al. [36] reported a strong dose-dependent GCaMP increase with 2.5 Hz and a milder offline inhibition at 20 Hz. The data analyzed in the present study comprised 5 on-target and 5 off-target animals at 2.5 Hz (210 on-target and 198 off-target trials total) and 5 on-target and 4 off-target animals at 20 Hz (216 on-target and 168 off-target trials total). Note that we excluded one animal from analysis (“cmtb3”) as it did not show any GCaMP response during or after stimulation (Supplementary Figs. S6–S7).

### 4.2 Preprocessing

Each trial was preprocessed independently using a three-step procedure. First, the isosbestic (405 nm) reference channel was regressed out of the GCaMP signal using ordinary least-squares regression *fit to the prestimulation baseline period only*, such that we did not inadvertently remove stimulation-evoked responses. The regression included an intercept term, and the fitted model was applied to the full trial to remove motion and hemodynamic artifacts common to both channels. Next, a linear trend was fit to the full trial-length residual and subtracted to remove slow drift. Finally, the detrended signal was z-scored using the mean and standard deviation of the prestimulation baseline period, yielding a baseline-normalized signal in units of standard deviations. No temporal smoothing was applied, preserving the full temporal structure of the signal for subsequent irreversibility analysis. Note that our preprocessing differs somewhat from that of Murphy et al. [36], which is relevant when comparing findings across studies.

### 4.3 Ordinal entropy production rate estimation

To quantify temporal irreversibility in the preprocessed calcium signal, we employed ordinal pattern analysis [1]. This approach maps a scalar time series into a sequence of discrete symbols by replacing each delay-embedded vector with its ordinal pattern (the rank ordering of its components). The distribution of ordinal patterns carries information about the temporal structure of the signal, and its asymmetry under time reversal provides a nonparametric estimator of the entropy production rate (EPR) [42, 53, 55].

Specifically, for a time series segment {*x_t_*} of length *N*, we constructed delay embeddings of dimension *m* and delay *τ*:

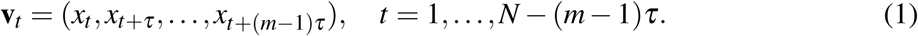

Each embedding vector **v***_t_* was mapped to one of *m*! possible ordinal patterns by recording the rank ordering of its components. The same procedure was applied to the time-reversed signal {*x_N_*_+1−*t*_}, yielding a backward ordinal pattern distribution. The symmetrized Kullback–Leibler (KL) divergence between the forward and backward distributions served as our EPR estimator:

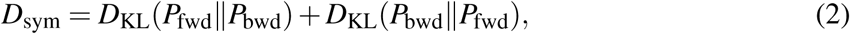

where *P*_fwd_ and *P*_bwd_ denote the normalized ordinal pattern frequency distributions of the forward and reversed signal, respectively, and where the KL divergence between two discrete distributions *P* and *Q* over the *m*! ordinal patterns *π_i_* is:

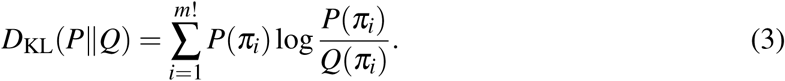

Additive Laplace smoothing (*ε* = 10^−6^ per bin) was applied before normalization to avoid undefined logarithms.

#### Symmetrized versus strict EPR estimator

The trajectory-level EPR is rigorously defined as the one-sided KL divergence between forward and time-reversed path distributions, *D*_fwd_ ≡ *D*_KL_(*P*_fwd_∥*P*_bwd_) [42, 47]. We adopted the symmetrized form *D*_sym_ (the Jeffreys divergence [23]) following the ordinal-pattern irreversibility literature [53, 55], which employs it due to its improved finite-sample stability: it spreads the cost of any zero-frequency pattern symmetrically across both directions, rather than placing it asymmetrically on whichever distribution happens to lack support at a given element. To verify that this choice does not influence our reported inferences, we re-ran the dose-response analysis of Figure 2 using *D*_fwd_ in place of *D*_sym_. The resulting irreversibility measures *D*_sym_ and *D*_fwd_ were essentially proportional (*r >* 0.99; median *D*_sym_*/D*_fwd_ ratio ≈ 2.0). This is expected from the small-divergence regime in which our ordinal-pattern estimates lie: to leading order in ∥*P*_fwd_ − *P*_bwd_∥, both KL directions reduce to the same functional, so *D*_KL_(*P*_fwd_∥*P*_bwd_) ≈ *D*_KL_(*P*_bwd_∥*P*_fwd_) and *D*_sym_ ≈ 2 *D*_fwd_ [10, Ch. 11]. We therefore retained *D*_sym_ as our irreversibility measure throughout.

#### Parameter selection

We employed *m* = 3 and *τ* = 8 samples as the parameters for measuring ordinal irreversibility. We fixed *m* = 3 to keep the ordinal alphabet at 6 states, which promotes stable estimates compared to higher-dimensional embeddings. The delay parameter *τ* was then selected outcome-blind from the set {2, 4, 6, 8} using a baseline-only reliability sweep at fixed *m* = 3. The delay value *τ* = 8 exhibited the highest split-half correlation for animal-level baseline EPR and was therefore adopted as the primary setting. Sensitivity analyses were conducted at neighbouring delay scales and with a higher-dimensional embedding to assess robustness to alternative choices (see Section 2.4).

#### Temporal windowing

Each trial was segmented into three temporal windows: a prestimulation baseline (5 s, the maximum prestimulation time recorded by Murphy et al. [36]), stimulation (5 seconds for 2.5 Hz PRF; 40 s for 20 Hz PRF), and post-stimulation recovery (10 s beginning immediately after sonication ends). A 0.25-second boundary trim was applied at the baseline–stimulation transition to avoid edge effects.

### 4.4 Statistical analysis

We employed both animal-and trial-level analyses depending on the question being addressed. For the dose-reponse and EPR versus GCaMP analyses of Figures 2, 3, and 4, irreversibility was estimated after pooling trials of each animal and forming one estimate per animal. To investigate the influence of baseline irreversibility on the subsequent response to stimulation (Figs. 5,6), EPR was estimated separately for each trial to afford greater statistical power, trading off sample size with the SNR of each individual estimate.

#### Pooled-trial EPR and calcium amplitude measures

For each (animal × intensity × condition × window) cell, we computed one *D*_sym_ value by pooling ordinal-pattern counts across all trials belonging to that cell and then evaluating Eq. 2 on the pooled distribution (see Section 4.3). For GCaMP, the dependent variable was the mean fluorescence amplitude averaged across trials within the same cell. The changes from baseline were defined as Δ*D*_sym_ = *D*_sym_^window^ − *D*_sym_^baseline^ and ΔGCaMP = GCaMP^window^ −GCaMP^baseline^ for EPR and calcium amplitude, respectively. We note that change scores were computed primarily for visualization of effects – when conducting formal inference (see next section), we opted for ANCOVA-style mixed models where baseline value was included as a covariate.

#### Mixed-effects models

For formal inference, we fit linear mixed-effects models (LMMs) with animal as a random intercept [25], using the same ANCOVA-style specification for both EPR and GCaMP:

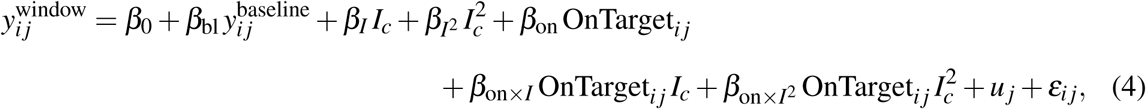

where *j* indexes the animal, *i* indexes the repeated measure (one sample per intensity × condition), *y* denotes either *D*_sym_ or mean GCaMP, the superscript “window” denotes the temporal window (stimulation or recovery), *I_c_* is the centered acoustic intensity, OnTarget*_i j_* is a binary indicator equal to 1 for on-target and 0 for off-target, and *u _j_* ∼ *N* (0*, σ* ^2^) is the random intercept for animal *j*. Including the baseline value as a covariate adjusts for variability in prestimulation state while avoiding the mechanical baseline–change coupling that arises from using difference scores as the outcome. The main-effect dose terms (*β_I_*, *β_I_*_2_) capture any shared dose dependence across conditions, while the interaction terms *β*_on×*I*_ and *β*_on×*I*2_ test whether the linear and quadratic dose dependence is specific to on-target stimulation. Models were fit using restricted maximum likeli-hood (REML) with the L-BFGS solver as implemented in the statsmodels Python package.

#### EPR beyond calcium amplitude

To test whether irreversibility carries dose information beyond what calcium amplitude can explain, we fit a hierarchy of three nested mixed-effects models on the pooled-trial data. All three shared a random intercept for animal.

Model A (baseline + amplitude only) included only prestimulation EPR and the window-matched fluorescence change as predictors:

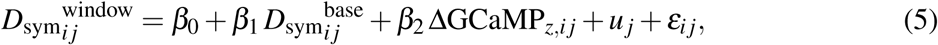

where ΔGCaMP*_z,i j_* is the standardized change in mean calcium amplitude for repeated measure *i* of animal *j*. Model B (shared dose) added linear and quadratic intensity terms that were common to both on-and off-target conditions:

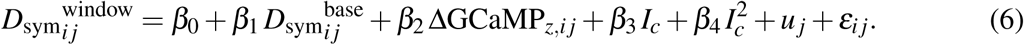

Model C (condition-specific dose) added an on-target fixed effect and further allowed the dose dependence to differ between on-target and off-target stimulation:

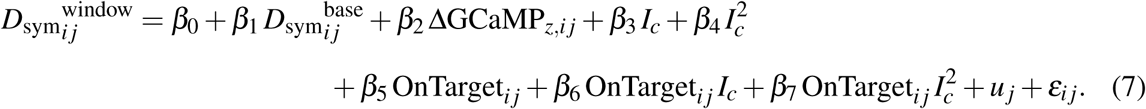

We employed likelihood-ratio (LR) tests to assess significant increases in explained variance: comparing Model B to Model A tests whether acoustic dose explains any EPR variance beyond baseline state and calcium amplitude change. Comparing Model C to Model B tests whether the dose–EPR relationship differs between on-target and off-target stimulation. Finally, to ask whether dose shapes EPR within the on-target condition alone, we refit Models A and B on the on-target subset only and compared them with a third LR test.

### 4.5 Baseline neural state and response prediction

To test whether *prestimulation* temporal irreversibility predicts the response to sonication, we measured baseline EPR at the level of individual trials, focusing the analysis on the 2.5 Hz PRF recordings.

#### Trial-level EPR and GCaMP computation

For each trial, we computed ordinal EPR and mean GCaMP amplitude separately for the baseline, stimulation, and recovery windows. Trial-level change scores were defined as Δ*D*_sym_ = *D*_sym_^window^ − *D*_sym_^baseline^ and ΔGCaMP = 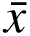^window^ − 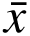^baseline^, where 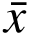 denotes the mean calcium amplitude within the specified window.

#### Baseline-state mixed-effects models

To test whether baseline state modulates the response to tFUS, we pooled trials across acoustic intensities and discretised baseline EPR into terciles (Low, Mid, High). For the EPR outcome measure, we fit a hierarchy of mixed-effects models according to:

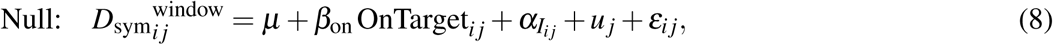

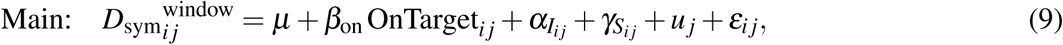

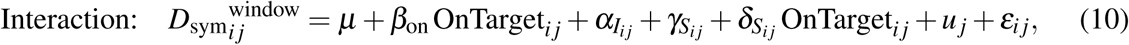

where *α_Ii j_* denotes fixed effects of acoustic intensity (entered as a categorical factor with six levels), *S_i j_* ∈ {Low, Mid, High} is the baseline-EPR tercile, *γ_Si j_* is the corresponding tercile fixed effect, *δ_Si j_* captures the tercile-by-condition interaction, and *u _j_* ∼ *N* (0*, σ* ^2^) is a random intercept for animal *j*. LR tests compared the Main model to the Null (baseline-state main effect) and the Interaction model to the Main (baseline-state × condition interaction).

The choice of a categorical form for intensity reflects the non-monotonic dose-response of EPR (Fig. 4B) – modeling intensity as categorical makes no assumption about the shape of the dose-response and thus avoids introducing misspecification into the estimate of the quantity-of-interest, namely the baseline-on-target interaction.

For GCaMP, the same three-model hierarchy was applied with calcium amplitude as the outcome and baseline GCaMP added as a covariate to adjust for prestimulation fluorescence:

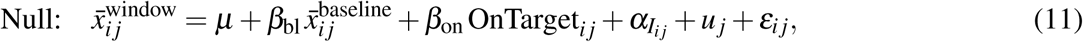

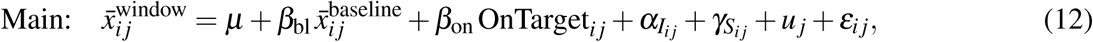

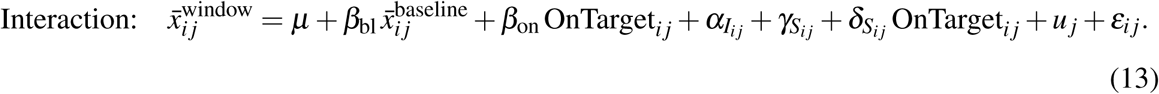

#### Incremental prediction of GCaMP by baseline EPR

To quantify whether baseline EPR car-ries predictive information about the subsequent GCaMP response beyond what baseline GCaMP and acoustic dose already explain, we fit trial-level mixed models on the on-target trials only. The reduced model predicted GCaMP from its baseline value, centred intensity (linear and quadratic), and a random animal intercept:

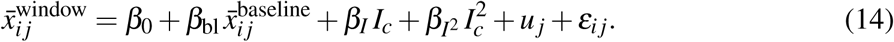

The full model included baseline *D*_sym_ as an additional predictor:

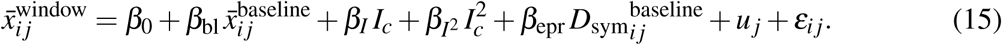

A significant LR test comparing the full model to the reduced model indicates that baseline EPR explains GCaMP variance that is not captured by baseline fluorescence or dose. Both models were fit with ML estimation. The added-variable (partial regression) plots in Figure 6A,B visualise the residual association between baseline EPR and post-window GCaMP after removing the shared covariates [18].

## Supplementary Material

**Figure S1:**
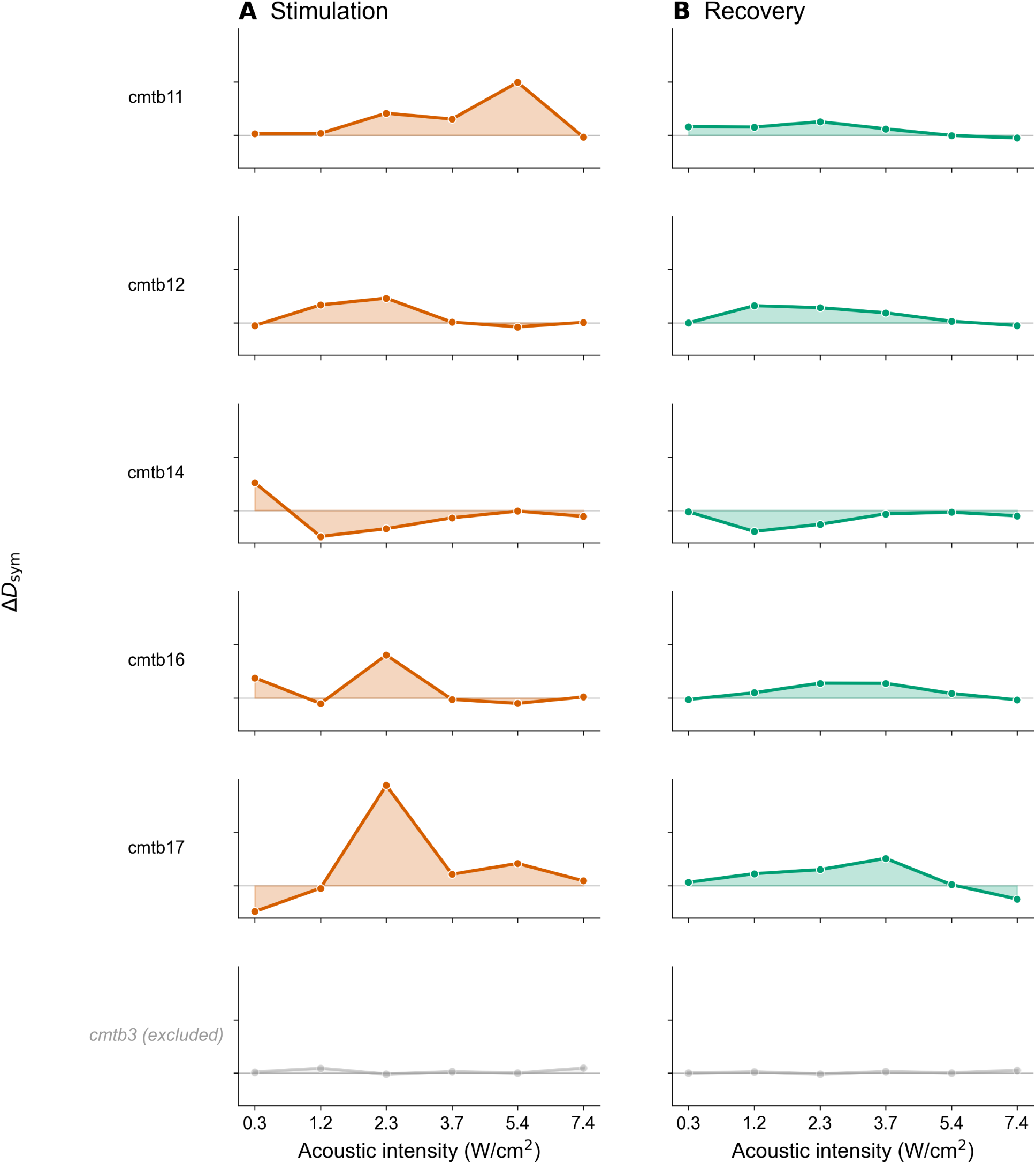
Per-animal. Δ*D*_sym_ **dose-response: 2.5 Hz on-target.** Each row shows one animal’s Δ*D*_sym_ as a function of acoustic intensity; left column: stimulation, right column: recovery. All panels share a common *y*-axis to facilitate comparison across animals. One animal (cmtb3, gray italic) was excluded from the primary analysis because it did not show any GCaMP response (see Supplementary Figs. S6–S7). Among the remaining 5 animals, a non-monotonic recovery profile peaking between 1.2 and 3.7 W/cm^2^ is visible in 4, indicating that the group-level quadratic effect is not driven by a single subject.

**Figure S2:**
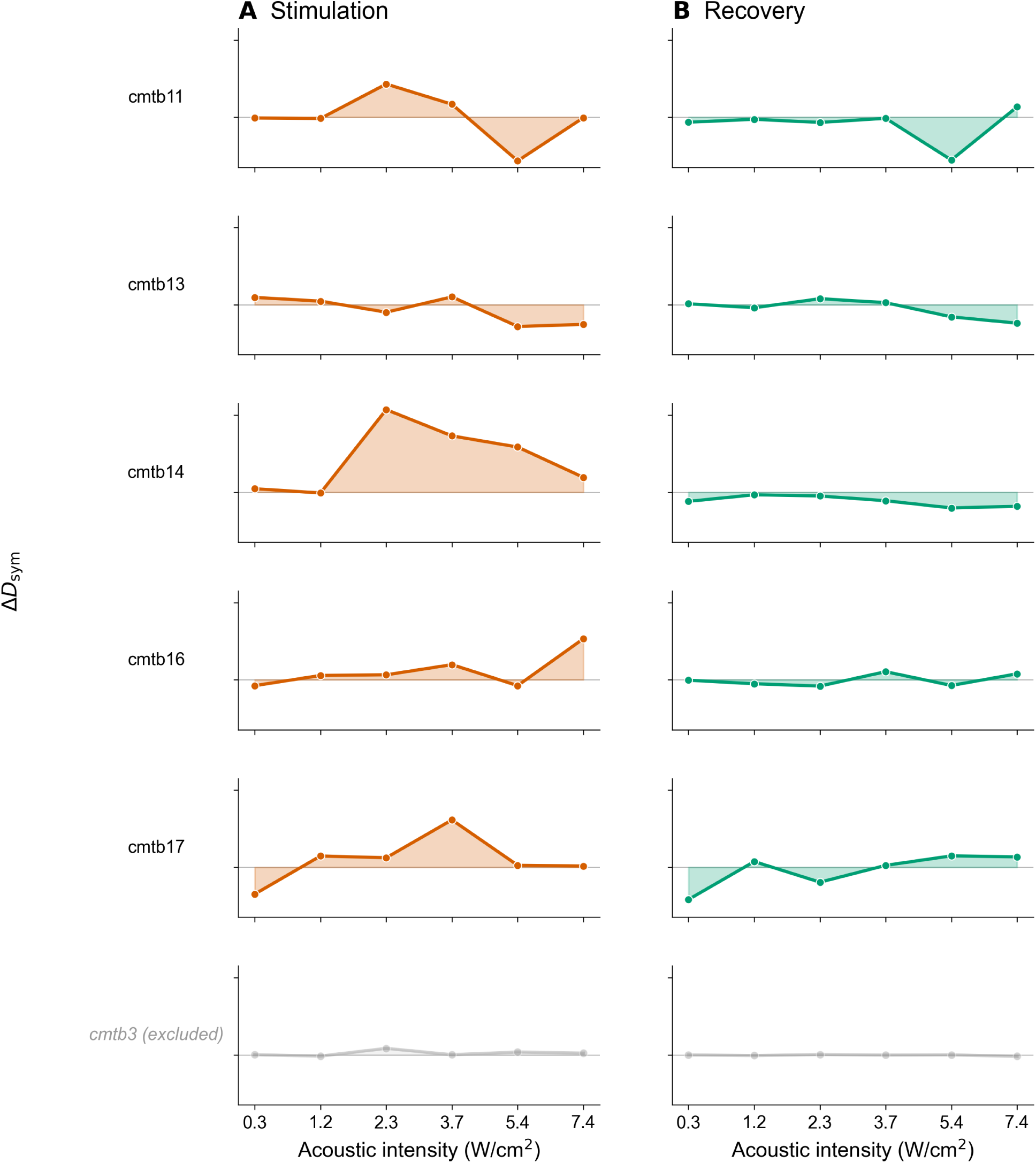
Per-animal. Δ*D*_sym_ **dose-response: 2.5 Hz off-target.** Same format as Fig. S1. Inter-estingly, off-target stimulation generates an inverted-U dose-response in 3 of 5 animals. During the recovery period, EPR changes fluctuate around zero with no systematic intensity dependence.

**Figure S3:**
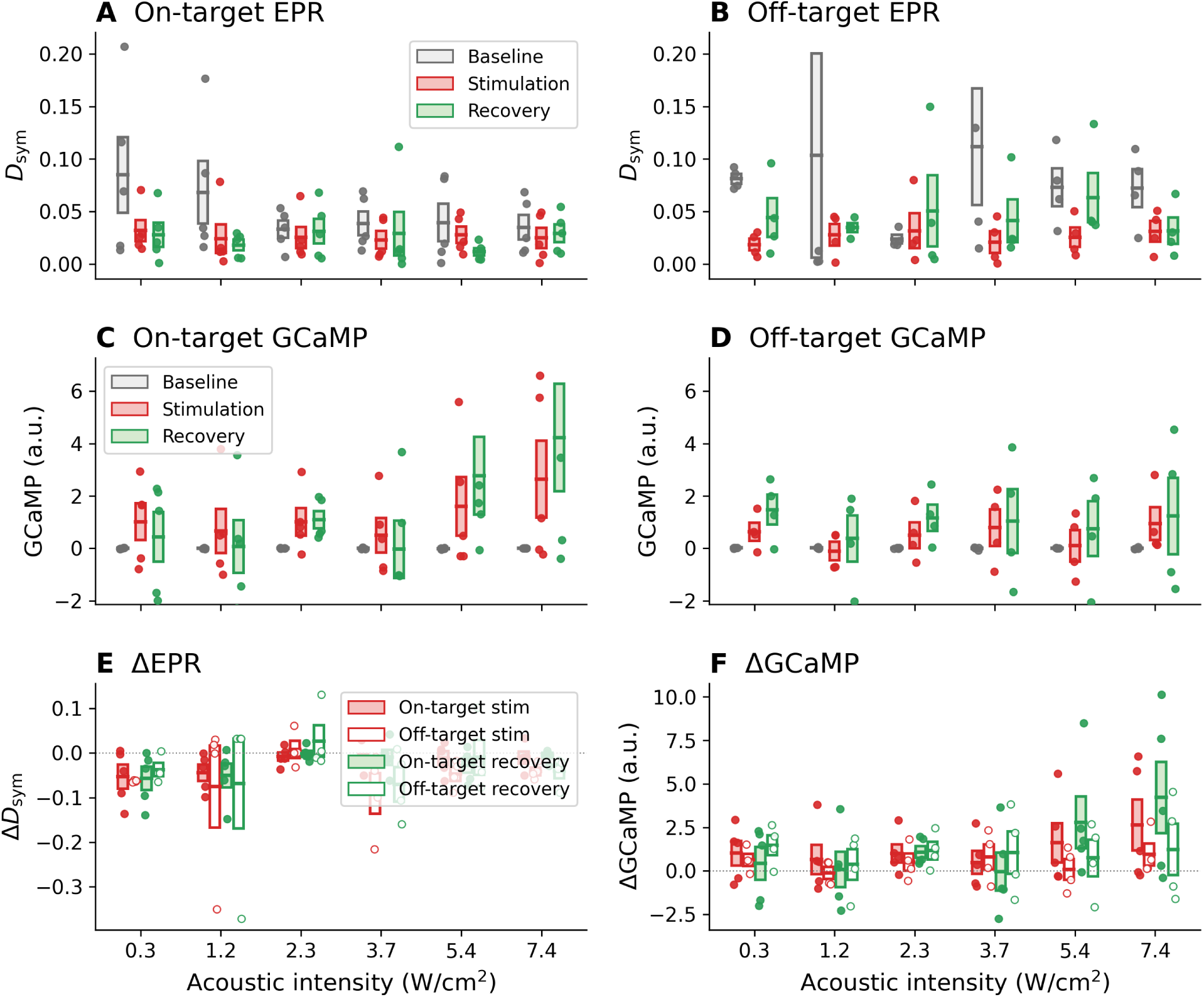
EPR and GCaMP dose-response at 20 Hz PRF. Same layout as Fig. 2 but for 20 Hz stimulation of the CMT. Murphy et al. [36] reported an offline inhibition of calcium amplitude at this configuration. We were unable to resolve a main effect of on-target stimulation on the GCaMP amplitude. Similarly, the corresponding analysis of EPR did not resolve a significant main effect of on-target sonication during stimulation (*p* = 0.940), although a nominally significant EPR *reduction* was present during recovery (*p* = 0.044).

**Figure S4:**
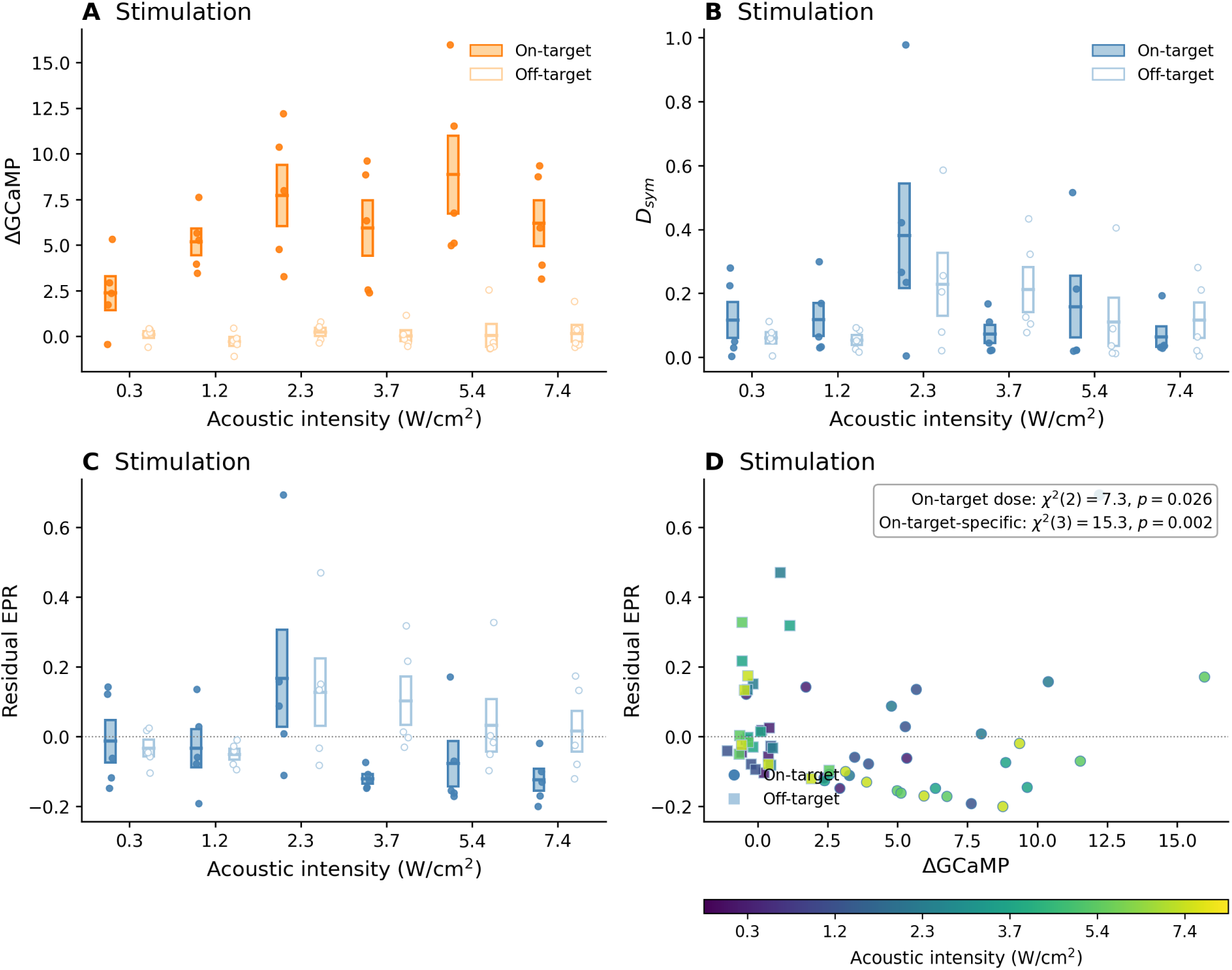
Stimulation-window EPR cannot be fully explained by calcium amplitude. Same analysis as Fig. 4 but applied to the stimulation window rather than recovery. **(A)** The change in GCaMP during sonication (ΔGCaMP) rises strongly with dose after on-target tFUS, while off-target remains near zero. **(B)** The EPR during sonication (*D*_sym_) shows condition-specific dose structure. **(C)** The residual EPR after regressing out ΔGCaMP and baseline EPR: on-target residual dose structure is present but weaker than in the recovery analysis (Fig. 4C). Error bars: ±1 SEM across animals. **(D)** Residual EPR versus ΔGCaMP, coloured by intensity. Within the on-target condition, dose remained significant after accounting for baseline EPR and ΔGCaMP (*χ*^2^(2) = 7.28, *p* = 0.026). Allowing dose terms to differ between on-target and off-target fur-ther improved fit (*χ*^2^(3) = 15.27, *p* = 0.002). Unlike recovery, however, the ΔGCaMP covariate remained strongly positive (*β* = +0.181, *p <* 0.001), indicating that stimulation-window EPR is only partially dissociated from fluorescence amplitude.

**Figure S5:**
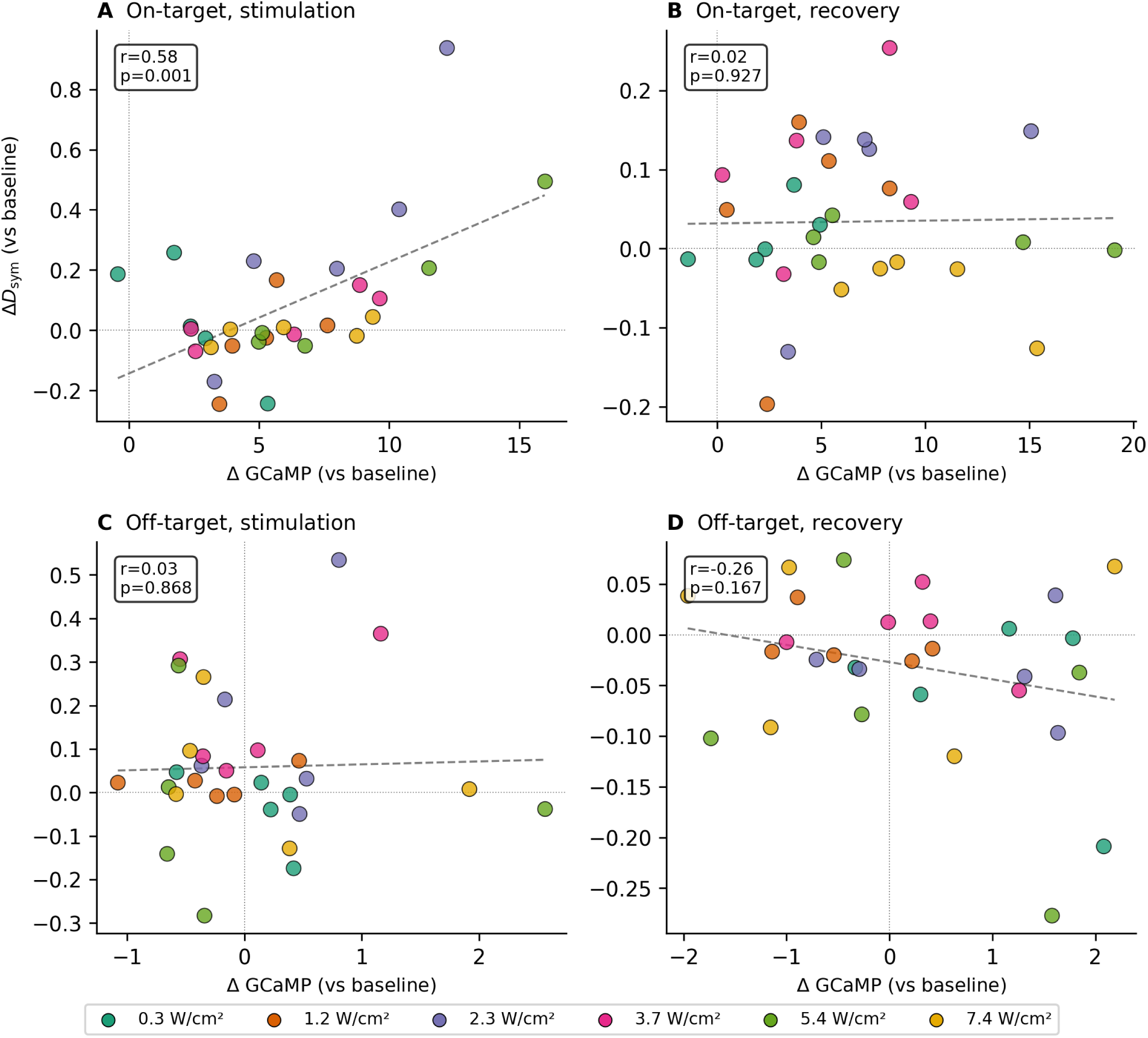
Correlation between. Δ*D*_sym_ **and** Δ**GCaMP across conditions and windows at 2.5 Hz.** Each point represents one animal×intensity combination; colours denote acoustic inten-sity. **(A)** On-target, stimulation: *r* = 0.58, *p <* 0.001. **(B)** On-target, recovery: *r* = 0.02, *p* = 0.927. **(C)** Off-target, stimulation: *r* = 0.03, *p* = 0.868. **(D)** Off-target, recovery: *r* = −0.26, *p* = 0.167. Main text Fig. 3C,D shows on-target panels only.

**Figure S6:**
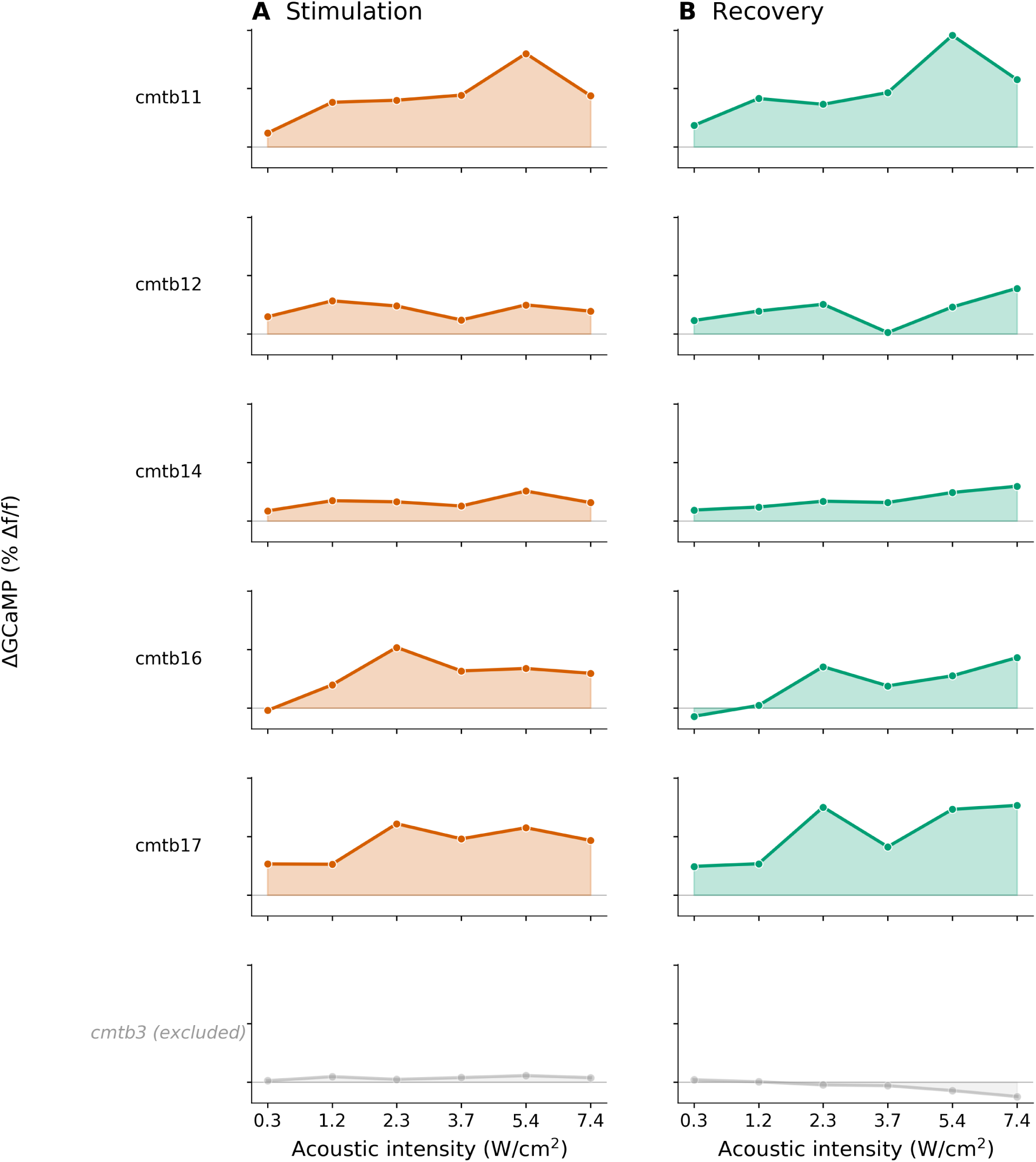
Per-animal. Δ**GCaMP dose–response: 2.5 Hz on-target.** Each row shows one ani-mal’s baseline-subtracted GCaMP amplitude as a function of acoustic intensity; left column: stim-ulation, right column: recovery. All panels share a common *y*-axis. One animal (cmtb3, gray italic) was excluded from analysis as it did not exhibit a calcium response during or after stimulation. Remaining animals show positive responses that increase with intensity, consistent with the monotonic group-level dose-response (Fig. 2).

**Figure S7:**
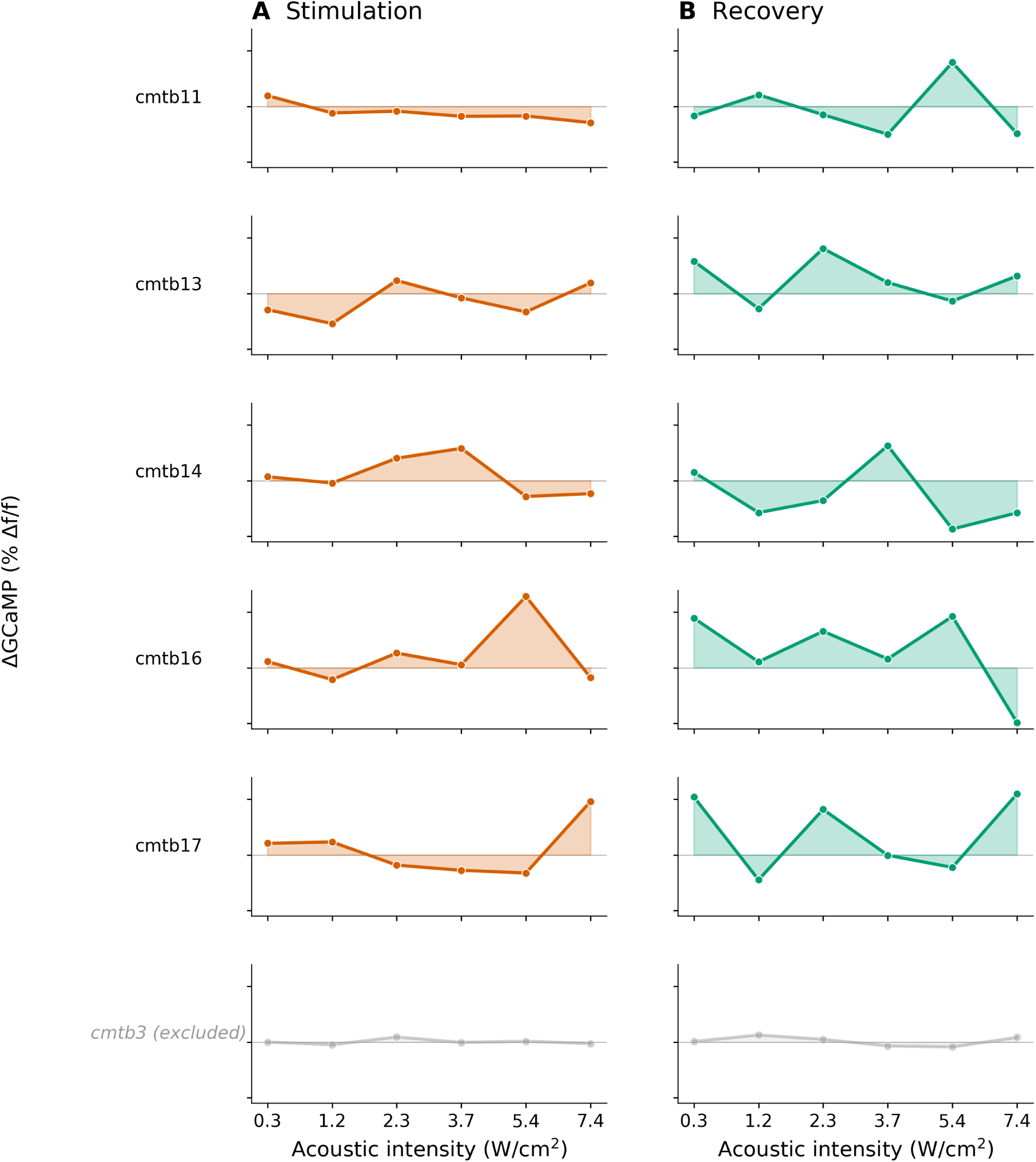
**Per-animal**Δ**GCaMP dose–response: 2.5 Hz off-target.** Same format as Fig. S6. Off-target changes remain near zero with no consistent dose dependence.

---

**Table S1:** Hierarchical mixed-model tests: EPR dose structure beyond baseline and calcium amplitude. Each row reports a likelihood-ratio (LR) test between nested models. Model A (base-line + amplitude only): *D*_sym_^post^ ∼ *D*_sym_^base^ + ΔGCaMP*_z_* + (1|animal). Model B (shared dose): adds *I_c_* + *Ic c*^2^. Model C (condition-specific dose): adds OnTarget + OnTarget×*I_c_* + OnTarget×*Ic c*^2^. “B vs. A (on-target)” tests dose structure in on-target trials only; “B vs. A (all)” pools on-target and off-target; “C vs. B” tests whether dose–EPR slopes differ between conditions. *β*_ΔGCaMP_ and its *p*-value are from Model C.

| Config. | Window | Comparison | $\chi^2$ | df | $p$ | $\beta_{\Delta\text{GCaMP}}$ | $p_{\Delta\text{GCaMP}}$ |
| --- | --- | --- | --- | --- | --- | --- | --- |
| *CMT 2.5 Hz (n=5 on-target, n=5 off-target)* |  |  |  |  |  |  |  |
|  | Stim | B vs. A (on-target) | 7.28 | 2 | .026 | +0.170 | .113 |
|  | Stim | B vs. A (all) | 3.87 | 2 | .144 | +0.181 | < .001 |
|  | Stim | C vs. B | 15.27 | 3 | .002 | +0.181 | < .001 |
|  | Rec | B vs. A (on-target) | 13.55 | 2 | .001 | −0.013 | .469 |
|  | Rec | B vs. A (all) | 7.23 | 2 | .027 | −0.006 | .642 |
|  | Rec | C vs. B | 27.23 | 3 | < .001 | −0.006 | .642 |
| *CMT 20 Hz (n=5 on-target, n=4 off-target)* |  |  |  |  |  |  |  |
|  | Stim | B vs. A (on-target) | 1.04 | 2 | .594 | +0.003 | .445 |
|  | Stim | C vs. B | 0.72 | 4 | .949 | +0.002 | .575 |
|  | Rec | B vs. A (on-target) | 0.44 | 2 | .804 | −0.003 | .319 |
|  | Rec | C vs. B | 0.49 | 4 | .974 | −0.001 | .780 |

## References

[1] C. Bandt and B. Pompe. Permutation entropy: a natural complexity measure for time series. Physical Review Letters, 88(17):174102, 2002. doi: 10.1103/PhysRevLett.88.174102.

[2] G. Batsikadze, V. Moliadze, W. Paulus, M.-F. Kuo, and M. A. Nitsche. Partially non-linear stimulation intensity-dependent effects of direct current stimulation on motor cortex excitability in humans. The Journal of Physiology, 591(7):1987–2000, 2013. doi: 10.1113/jphysiol.2012.249730.

[3] C. Battle, C. P. Broedersz, N. Fakhri, V. F. Geyer, J. Howard, C. F. Schmidt, and F. C. MacKintosh. Broken detailed balance at mesoscopic scales in active biological systems. Science, 352(6285):604–607, 2016. doi: 10.1126/science.aac8167.

[4] T. O. Bergmann, A. Karabanov, G. Hartwigsen, A. Thielscher, and H. R. Siebner. Combining non-invasive transcranial brain stimulation with neuroimaging and electrophysiology: Current approaches and future perspectives. NeuroImage, 140:4–19, 2016. doi: 10.1016/j.neuroimage.2016.02.012.

[5] J. Blackmore, S. Shrivastava, J. Sallet, C. R. Butler, and R. O. Cleveland. Ultrasound neuromodulation: A review of results, mechanisms and safety. Ultrasound in Medicine & Biology, 45(7):1509–1536, 2019. doi: 10.1016/j.ultrasmedbio.2018.12.015.

[6] A. Bystritsky, A. S. Korb, P. K. Douglas, M. S. Cohen, W. P. Melega, A. P. Mulgaonkar, A. DeSalles, B.-K. Min, and S.-S. Yoo. A review of low-intensity focused ultrasound pulsation. Brain Stimulation, 4(3):125–136, 2011. doi: 10.1016/j.brs.2011.03.007.

[7] H. Caffaratti, B. Slater, N. Shaheen, A. Rhone, R. Calmus, M. Kritikos, S. Kumar, B. Dlouhy, H. Oya, T. Griffiths, A. D. Boes, N. Trapp, M. Kaiser, J. Sallet, M. I. Banks, M. A. Howard, M. Zanaty, and C. I. Petkov. Neuromodulation with ultrasound: Hypotheses on the directionality of effects and community resource. medRxiv, 2025. doi: 10.1101/2024.06.14.24308829.

[8] E. J. Calabrese and L. A. Baldwin. Hormesis: the dose-response revolution. Annual Review of Pharmacology and Toxicology, 43:175–197, 2003. doi: 10.1146/annurev.pharmtox.43.100901.140223.

[9] T.-W. Chen, T. J. Wardill, Y. Sun, S. R. Pulver, S. L. Renninger, A. Baohan, E. R. Schreiter, R. A. Kerr, M. B. Orger, V. Jayaraman, et al. Ultrasensitive fluorescent proteins for imaging neuronal activity. Nature, 499(7458):295–300, 2013. doi: 10.1038/nature12354.

[10] T. M. Cover and J. A. Thomas. Elements of Information Theory. Wiley-Interscience, 2 edition, 2006. ISBN 0471241954.

[11] G. E. Crooks. Entropy production fluctuation theorem and the nonequilibrium work relation for free energy differences. Physical Review E, 60(3):2721–2726, 1999. doi: 10.1103/PhysRevE.60.2721.

[12] G. Deco, Y. Sanz Perl, H. Bocaccio, E. Tagliazucchi, and M. L. Kringelbach. The INSIDE-OUT framework provides precise signatures of the balance of intrinsic and extrinsic dynamics in brain states. Communications Biology, 5(1):578, 2022. doi: 10.1038/s42003-022-03505-7.

[13] S. Deffner and S. Campbell. Quantum thermodynamics: An introduction to the thermodynamics of quantum information. Morgan & Claypool Publishers, 2019. doi: 10.1088/2053-2571/ab21c6.

[14] J. Dell’Italia. Current state of potential mechanisms supporting low intensity focused ultrasound for neuromodulation. Frontiers in Human Neuroscience, 16:872639, 2022. doi: 10.3389/fnhum.2022.872639.

[15] Z. Esmaeilpour, P. Marangolo, B. M. Hampstead, S. Bestmann, E. Galletta, H. Knotkova, and M. Bikson. Incomplete evidence that increasing current intensity of tDCS boosts outcomes. Brain Stimulation, 11(2):310–321, 2018. doi: 10.1016/j.brs.2017.12.002.

[16] M. Esposito and C. Van den Broeck. Three faces of the second law. I. Master equation formulation. Physical Review E, 82(1):011143, 2010. doi: 10.1103/PhysRevE.82.011143.

[17] A. Fomenko, C. Neudorfer, R. F. Dallapiazza, S. K. Kalia, and A. M. Lozano. Low-intensity ultrasound neuromodulation: An overview of mechanisms and emerging human applications. Brain Stimulation, 11(6):1209–1217, 2018. doi: 10.1016/j.brs.2018.08.013.

[18] J. Fox. Applied Regression Analysis and Generalized Linear Models. Sage, Los Angeles, 3 edition, 2016.

[19] L. Gammaitoni, P. Hänggi, P. Jung, and F. Marchesoni. Stochastic resonance. Reviews of Modern Physics, 70(1):223–287, 1998. doi: 10.1103/RevModPhys.70.223.

[20] F. S. Gnesotto, F. Mura, J. Gladrow, and C. P. Broedersz. Broken detailed balance and non-equilibrium dynamics in living systems: a review. Reports on Progress in Physics, 81(6): 066601, 2018. doi: 10.1088/1361-6633/aab3ed.

[21] C. S. Herrmann, M. M. Murray, S. Ionta, A. Hutt, and J. Lefebvre. Shaping intrinsic neural oscillations with periodic stimulation. Journal of Neuroscience, 36(21):5328–5337, 2016. doi: 10.1523/JNEUROSCI.0236-16.2016.

[22] C. Jarzynski. Nonequilibrium equality for free energy differences. Physical Review Letters, 78(14):2690–2693, 1997. doi: 10.1103/PhysRevLett.78.2690.

[23] H. Jeffreys. An invariant form for the prior probability in estimation problems. Proceedings of the Royal Society A, 186(1007):453–461, 1946. doi: 10.1098/rspa.1946.0056.

[24] L. Lacasa, A. Núñez, É. Roldán, J. M. Parrondo, and B. Luque. Time series irreversibility: a visibility graph approach. The European Physical Journal B, 85(6):217, 2012. doi: 10.1140/epjb/e2012-20809-8.

[25] N. M. Laird and J. H. Ware. Random-effects models for longitudinal data. Biometrics, pages 963–974, 1982. doi: 10.2307/2529876.

[26] A. A. Legaria, B. A. Matikainen-Ankney, B. Yang, B. Ahanonu, J. A. Licholai, J. G. Parker, A. Bharioke, and A. V. Kravitz. Fiber photometry in striatum reflects primarily nonsomatic changes in calcium. Nature Neuroscience, 25(9):1124–1128, 2022. doi: 10.1038/s41593-022-01152-z.

[27] W. Legon, T. F. Sato, A. Opitz, J. Mueller, A. Barbour, A. Williams, and W. J. Tyler. Transcranial focused ultrasound modulates the activity of primary somatosensory cortex in humans. Nature Neuroscience, 17(2):322–329, 2014. doi: 10.1038/nn.3620.

[28] J. Li, J. M. Horowitz, T. R. Gingrich, and N. Fakhri. Quantifying dissipation using fluctuating currents. Nature Communications, 10(1):1666, 2019. doi: 10.1038/s41467-019-09631-x.

[29] S. Little, E. Tripoliti, M. Beudel, A. Pogosyan, H. Cagnan, D. Herz, S. Bestmann, T. Aziz, et al. Adaptive deep brain stimulation in advanced Parkinson disease. Annals of Neurology, 74(3):449–457, 2013. doi: 10.1002/ana.23951.

[30] D. Lucente, G. Gradenigo, and L. Salasnich. Entropy production and irreversibility in the linearized stochastic amari neural model. Entropy, 27(11):1104, 2025.

[31] C. W. Lynn, E. J. Cornblath, L. Papadopoulos, M. A. Bertolero, and D. S. Bassett. Broken detailed balance and entropy production in the human brain. Proceedings of the National Academy of Sciences, 118(47):e2109889118, 2021. doi: 10.1073/pnas.2109889118.

[32] C. W. Lynn, C. M. Holmes, W. Bialek, and D. J. Schwab. Decomposing the local arrow of time in interacting systems. Physical Review Letters, 129(11):118101, 2022. doi: 10.1103/PhysRevLett.129.118101.

[33] R. G. Mair, K. D. Onos, and J. R. Hembrook. Cognitive activation by central thalamic stimulation: The Yerkes-Dodson law revisited. Dose-Response, 9(3):313–331, 2011. doi: 10.2203/dose-response.10-017.Mair.

[34] I. A. Martínez, G. Bisker, J. M. Horowitz, and J. M. Parrondo. Inferring broken detailed balance in the absence of observable currents. Nature Communications, 10(1):3542, 2019. doi: 10.1038/s41467-019-11051-w.

[35] M. D. McDonnell and L. M. Ward. The benefits of noise in neural systems: bridging theory and experiment. Nature Reviews Neuroscience, 12(7):415–426, 2011. doi: 10.1038/nrn3061.

[36] K. R. Murphy, J. S. Farrell, J. Bendig, A. Mitra, C. Luff, I. A. Stelzer, H. Yamaguchi, C. C. Angelakos, M. Choi, W. Bian, T. DiIanni, E. M. Pujol, N. Matosevich, R. Airan, B. Gaudillière, E. E. Konofagou, K. Butts-Pauly, I. Soltesz, and L. de Lecea. Optimized ultrasound neuromodulation for non-invasive control of behavior and physiology. Neuron, 112(19): 3252–3266.e5, 2024. doi: 10.1016/j.neuron.2024.07.002.

[37] T. Nandi, B. R. Kop, K. Naftchi-Ardebili, C. J. Stagg, K. B. Pauly, and L. Verhagen. Biophysical effects and neuromodulatory dose of transcranial ultrasonic stimulation. ArXiv, 2024. Preprint: 2406.19869.

[38] O. Naor, S. Krupa, S. Bhatt, and D. K. Bhatt. Ultrasonic neuromodulation. Journal of Neural Engineering, 13(3):031003, 2016. doi: 10.1088/1741-2560/13/3/031003.

[39] D. T. Nguyen, D. E. Berisha, E. E. Konofagou, and J. P. Dmochowski. Neuronal responses to focused ultrasound are gated by pre-stimulation brain rhythms. Brain Stimulation, 15(1): 233–243, 2022. doi: 10.1016/j.brs.2022.01.002.

[40] G. Paxinos and K. B. J. Franklin. The Mouse Brain in Stereotaxic Coordinates. Academic Press, San Diego, 5 edition, 2019.

[41] A. Pikovsky, M. Rosenblum, and J. Kurths. Synchronization: A Universal Concept in Non-linear Sciences. Cambridge University Press, Cambridge, 2001.

[42] É. Roldán and J. M. Parrondo. Estimating dissipation from single stationary trajectories. Physical Review Letters, 105(15):150607, 2010. doi: 10.1103/PhysRevLett.105.150607.

[43] É. Roldán and J. M. Parrondo. Entropy production and Kullback-Leibler divergence between stationary trajectories of discrete systems. Physical Review E, 85(3):031129, 2012. doi: 10.1103/PhysRevE.85.031129.

[44] V. Romei, V. Brodbeck, C. Michel, A. Amedi, A. Pascual-Leone, and G. Thut. Spontaneous fluctuations in posterior *α*-band EEG activity reflect variability in excitability of human visual areas. Cerebral Cortex, 18(9):2010–2018, 2008. doi: 10.1093/cercor/bhm229.

[45] Y. Sanz Perl, C. Pallavicini, I. Pérez Ipiña, A. Demertzi, V. Bonhomme, M. Charlotte, C. Duclos, S. Laureys, and E. Tagliazucchi. Nonequilibrium brain dynamics as a signature of consciousness. Physical Review E, 104(1):014411, 2021. doi: 10.1103/PhysRevE.104.014411.

[46] A. Seif, M. Hafezi, and C. Jarzynski. Machine learning the thermodynamic arrow of time. Nature Physics, 17:105–113, 2021. doi: 10.1038/s41567-020-1018-2.

[47] U. Seifert. Stochastic thermodynamics, fluctuation theorems and molecular machines. Reports on Progress in Physics, 75(12):126001, 2012. doi: 10.1088/0034-4885/75/12/126001.

[48] J. Silvanto and Z. Cattaneo. Common framework for “virtual lesion” and state-dependent TMS: The facilitatory/suppressive range model of online TMS effects on behavior. Brain and Cognition, 119:32–38, 2017. doi: 10.1016/j.bandc.2017.09.007.

[49] Y. Tufail, A. Matyushov, N. Baldwin, M. L. Tauchmann, J. Georges, A. Yoshihiro, S. I. H. Tillery, and W. J. Tyler. Transcranial pulsed ultrasound stimulates intact brain circuits. Neuron, 66(5):681–694, 2010. doi: 10.1016/j.neuron.2010.05.008.

[50] W. J. Tyler, Y. Tufail, M. Finsterwald, M. L. Tauchmann, E. J. Olson, and C. Majestic. Remote excitation of neuronal circuits using low-intensity, low-frequency ultrasound. PLoS ONE, 3 (10):e3511, 2008. doi: 10.1371/journal.pone.0003511.

[51] P.-F. Yang, M. A. Phipps, A. T. Newton, S. Jonathan, T. J. Manuel, J. C. Gore, W. A. Grissom, C. F. Caskey, and L. M. Chen. Differential dose responses of transcranial focused ultrasound at brain regions indicate causal interactions. Brain Stimulation, 15(6):1552–1564, 2022. doi: 10.1016/j.brs.2022.12.003.

[52] R. M. Yerkes and J. D. Dodson. The relation of strength of stimulus to rapidity of habit-formation. Journal of Comparative Neurology and Psychology, 18(5):459–482, 1908. doi: 10.1002/cne.920180503.

[53] M. Zanin. Ordinal patterns-based methodologies for distinguishing chaos from noise in discrete time series. Communications Physics, 4(1):190, 2021. doi: 10.1038/s42005-021-00696-z.

[54] M. Zanin and D. Papo. Algorithmic approaches for assessing irreversibility in time series: Review and comparison. Entropy, 23(11):1474, 2021. doi: 10.3390/e23111474.

[55] M. Zanin, A. Rodríguez-González, E. Menasalvas Ruiz, and D. Papo. Assessing time series reversibility through permutation patterns. Entropy, 20(9):665, 2018. doi: 10.3390/e20090665.

[56] T. Zhang, N. Pan, Y. Wang, C. Liu, and S. Hu. Transcranial focused ultrasound neuromodulation: A review of the excitatory and inhibitory effects on brain activity in human and animals. Frontiers in Human Neuroscience, 15:749162, 2021. doi: 10.3389/fnhum.2021.749162.

[57] B. Zrenner, C. Zrenner, P. C. Gordon, P. Belardinelli, E. J. McDermott, S. R. Soekadar, A. J. Fallgatter, U. Ziemann, and F. Müller-Dahlhaus. Brain oscillation-synchronized stimulation of the left dorsolateral prefrontal cortex in depression using real-time EEG-triggered TMS. Brain Stimulation, 13(1):197–205, 2020. doi: 10.1016/j.brs.2019.10.007.

[58] C. Zrenner, D. Desideri, P. Belardinelli, and U. Ziemann. Real-time EEG-defined excitability states determine efficacy of TMS-induced plasticity in human motor cortex. Brain Stimulation, 11(2):374–389, 2018. doi: 10.1016/j.brs.2017.11.016.

